# Allosteric Control and Glycan Shielding Adaptations in the SARS-CoV-2 Spike from Early to Peak Virulence

**DOI:** 10.1101/2025.03.11.642723

**Authors:** Srirupa Chakraborty, Kien N. Nguyen, Mingfei Zhao, S. Gnanakaran

## Abstract

The SARS-CoV-2 Spike glycoprotein is central to viral infectivity and immune evasion, making it a key target for vaccine and therapeutic design. This trimeric peplomer undergoes dynamic conformational changes, particularly in its Receptor Binding Domain (RBD), which transitions between closed (down) and ACE2-accessible (up) states relative to the rest of the protein, to facilitate host cell entry. Structural understanding of such critical inter-domain motions, as well as epitope exposure quantification, is essential for obtaining an effective molecular handle over this protein and, in turn, exploiting it towards improved immunogen development. Focusing on the early circulating D614G form and the later emerging Delta (B.1.617.2) variant with higher virulence, we performed large-scale molecular dynamics simulations of the soluble form of the Spike in both ‘down’ and ‘up’ conformations of the RBD. Guided by differences in overall fluctuations, we described reaction coordinates based on domain rotations and tilting to extract features that distinguish D614G versus Delta structural behavior of the N-terminal Domain (NTD) and RBD. Using reaction coordinate analysis and Principal Component Analysis (PCA), we identify allosteric coupling between the N-terminal Domain (NTD) and RBD, where NTD tilting influences RBD gating. While some of these motions are conserved across variants, Delta exhibits an optimized RBD-gating mechanism that enhances ACE2 accessibility. Additionally, glycan remodeling in Delta enhances shielding at the NTD supersite, contributing to reduced sensitivity to neutralizing antibodies. Finally, we uncover the impact of the D950N mutation in the HR1 region, which modulates downstream Spike dynamics and immune evasion. Together, our findings reveal variant-specific and conserved structural determinants of SARS-CoV-2 Spike function, providing a mechanistic basis for allosteric modulation, glycan-mediated immune evasion, and viral adaptation. These insights offer valuable guidance for rational vaccine and therapeutic design against SARS-CoV-2 and emerging variants.

## INTRODUCTION

Viruses, more notably RNA viruses, constantly undergo mutations that can evolve into new variants and strains[1, 2]. While most of these variants have minimal impact and disappear quickly, others can affect transmissibility, virulence, and vaccine effectiveness, persisting as variants of concern through natural selection. Since its discovery in December 2019, the Severe Acute Respiratory Syndrome Coronavirus 2 (SARS CoV-2) has accumulated several mutations. Based on the WHO epidemiological updates, SARS CoV-2 went through many Variants of Concern (such as the Alpha, Beta, Gamma, Delta, and Omicron lineages), along with several Variants of Interest for the SARS CoV-2[3, 4]. While Omicron out-competed other previous variants in global dissemination through its high transmissibility[5, 6], the Delta variant exhibited higher virulence, symptom severity, increased hospitalization, and long-COVID associations[7–9].

A large fraction of these mutations occur on the Spike peplomers expressed at the surface of the virus. This glycoprotein, which aids in host cell recognition, fusion, and cellular ingress[10], is the primary exposed viral antigen and is critical for triggering varieties of host immune responses[11]. Inversely, the Spike protein itself participates in immune evasion through glycan shielding and garnering mutations. Understanding the structure and dynamics of this protein and the effects of the emerging mutations is critical to improving immunogen design and treatment strategies that target viral entry.

The SARS CoV-2 Spike protein is a homotrimer, each of which accommodates two bulky and dynamic globular domains at the apical region – the N-terminal Domain (NTD) and the Receptor Binding Domain (RBD) (**Fig 1**). Both these domains have been found to prompt a large panel of antibodies, housing some critical epitopes[12, 13]. This manifestly indicates that obstructing the dynamics and functions of these domains can prevent the overall functioning of the virus. While it is known that the RBD transitions between ACE2-inaccessible “down” and ACE2-binding compatible “up” conformations[14], along with insights into some of the structural determinants of these transitions[15, 16], broader allosteric regulation of RBD dynamics, particularly its coupling with the NTD and other Spike domains, remains incompletely understood. The NTD has remained even more cryptic in terms of its structure-function relations. The relatively narrow base of the Spike has the furin cleavage site (FCS) with the RRAR motif on each protomer that separates the S1-S2 subunits[17]. The base also includes the heptad repeat helices HR1 and HR2, typical of class I fusion proteins.

**Figure 1.**
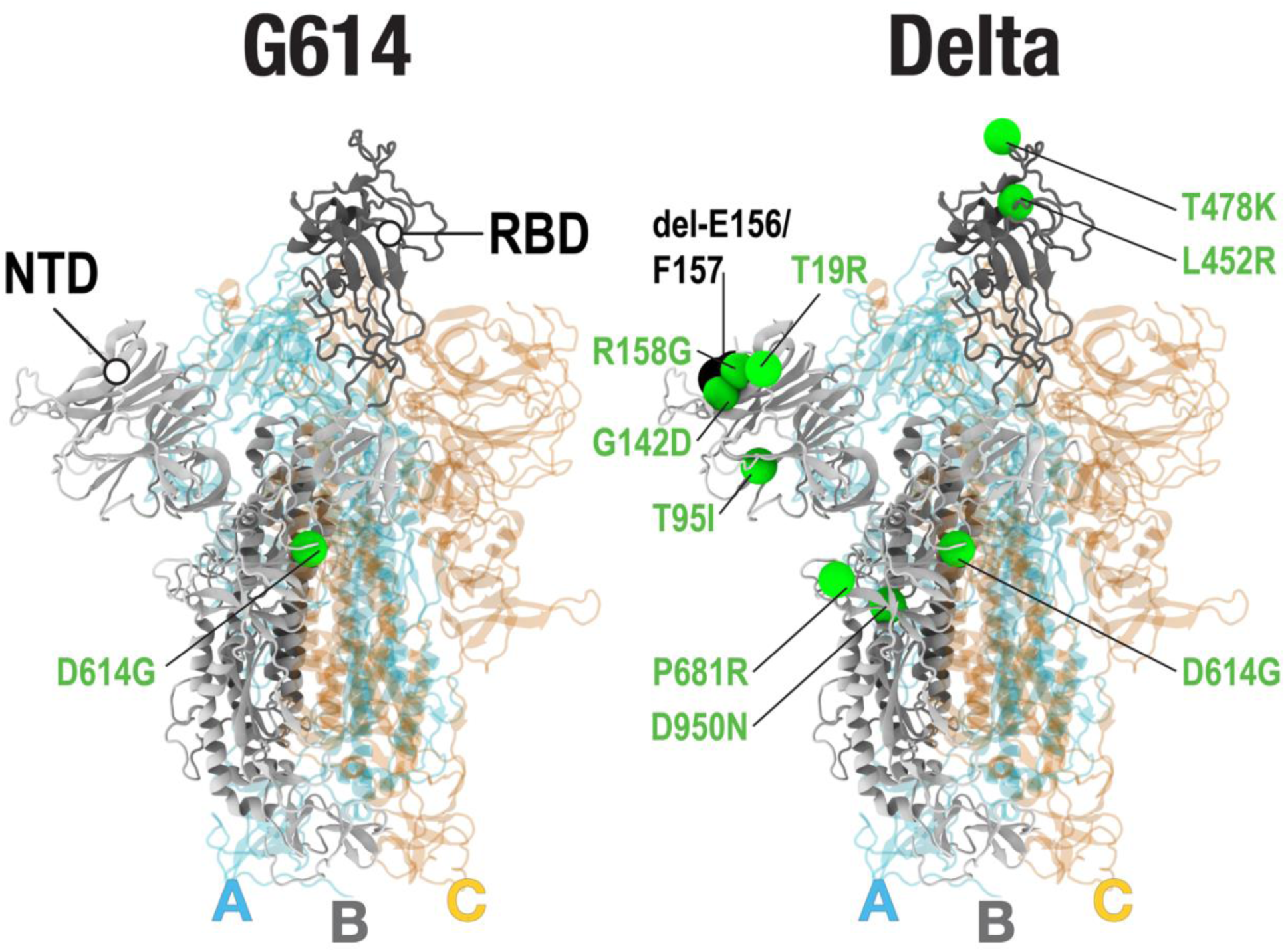
Structural description of the Spike protein, with the mutation/deletion sites that define the G614 and Delta variants. Point mutations are represented by green beads, and residue deletions by black beads. (Left) Depiction of the G614 Spike variant with the D614G mutation. (Right) Depiction of the Delta Spike variant. The NTD of Delta contains four mutations (T19R, T95I, G142D, and R158G), and two deletions (del-E156 and del-F157). The RBD of Delta harbors two mutations, L452R and T478K. Additional mutations in Delta are D614G, P681R, and D950N. In each Spike, the protomers “A, B and C” are colored cyan, gray and orange, respectively. In each spike, the RBD of protomer B is in the up conformation, while the RBDs of protomers A and C are in the down conformation. All molecular images were created using VMD[63].

A lot of experimental efforts have been expended to obtain details of the Spike protein structure through numerous studies[14, 18–22]. These, however, only manage to capture the static conformations of the protein in a small number of stable states. These proteins are dynamic entities, where not only the large-scale transitional motions but even the equilibrium fluctuations are critical elements that drive function. Exemplifying this are the effects of some emergent mutations over the last few years. While many of these sequence modifications can be easily interpreted by local structural changes[20, 22–25] and attributed to direct interaction outcomes (antibody escape or enhanced receptor affinity)[26–28], some important mutations are located away from known epitopes and ACE2 binding site. These mutations do not cause sizeable structural modifications, yet affect the Spike behavior through substantial alterations to essential dynamics and allosteric effects. A notable example is the single amino acid substitution D614G[29] located on an unstructured loop, away from key structural domains such as NTD, RBD, FCS and the HRs. Yet, by a cascade of structural dynamics, the D614G causes a higher probability of RBD opening[30, 31] and enhanced infection capability[29, 32], resulting in this substitution being the dominant variant.

The collective motions of NTD and RBD in SARS-CoV-2 variants have drawn interest due to their regulatory role in the functional change of Spike and potential drug-binding ability.[24, 33–36] Melero et al. have characterized NTD-RBD movements to a large degree as a rotation using a consensus-based image-processing approach with principal component analysis, but they also identified an important structural rearrangement from a complex pattern of flexibility presented by different RBDs.[35] Using molecular dynamics simulations, Li et al. found the neighboring NTD prohibits the RBD’s movements as a wedge detaches and swings away, leading to the potential binding target of NTD-RBD interface.[34] Choi et al. also used long all-atom simulations to study motions of the RBD and the NTD in fully glycosylated systems and non-glycosylated systems with full-length Spike in a viral membrane.[25] Wang et al. investigated conformational change of Delta Spike protein using molecular simulations to capture a “breath” motion initiated by an RBD swing movement, in which the three inter-protomer NTD-RBD pairs tilt outward and downward simultaneously.[24]

While numerous SARS-CoV-2 variants emerged over the course of the pandemic, the Delta variant is a crucial model for understanding viral evolution, structural adaptability, and immune evasion strategies. Unlike later variants, such as Omicron, which prioritized increased transmissibility at the cost of virulence, Delta exhibited a uniquely severe disease phenotype, contributing to higher hospitalization rates and mortality compared to both its predecessors and successors[8]. Notably, Delta’s defining mutations, including L452R, T478K, and P681R, enhanced its infectivity, RBD dynamics, and immune escape mechanisms, some of which persist in later Omicron sublineages[37]. Through its impact, Delta can serve as a benchmark for studying allosteric regulation, glycan-mediated shielding, and structural plasticity in SARS-CoV-2, providing insights that extend beyond a single variant and inform future vaccine and therapeutic design. The Delta (B.1.617.2) variant has 10 mutations on the Spike, over and above the D614G variant as depicted in **Fig. 1** — T19R,T95I,G142D,E156del,F157del and R158G in the NTD, L452R,T478K in the RBD, P681R near the FCS, and D950N at the base of the HR1 helix. Another salient feature of the Spike is that there are ∼22 N-glycans (as well as a few putative O-glycans)[38, 39] present on the surface of the protein that provide immune shielding and operational controls by virtue of their dynamics[40–42], which cannot be fully resolved by current experimental techniques. To better understand the functional nuances of these mutations and glycans, computational modeling of the detailed dynamics of this glycoprotein is the way forward. Findings from such simulations can provide a thorough understanding of the effects of emerging mutations as well as the innate dynamic properties that should remain invariant.

In this article, we investigate how the inter-domain dynamics between RBD and NTD vary between two different variants, Delta and the D614G (called G614 from here on). The latter variant is one of the major early substitutions on the original Wuhan strain, which was widely probed in terms of structure-dynamics relationships[20, 30, 40, 43]. To dissect how mutations can affect Spike function as well as extract the innate structural invariants that drive the peplomer behavior, we performed extensive all-atom explicit-solvent molecular dynamics (MD) simulations of the G614 and Delta Spike. For each of these variants, the ‘1-up’ and ‘all-down’ conformations (indicating the up/down orientation of RBD relative to the Spike) were modeled with native glycosylation patterns[39]. We begin by studying the glycan shielding of the Spike protein surface and how the virus evolved to optimize the shielding effect over critical regions. Guided by differences in overall fluctuations, we further define cardinal reaction coordinates based on NTD/RBD rotations and tilting to extract features that distinguish variant-dependent structural behavior and consequently establish the dynamic links that drive concerted motions between these two important domains. We go on to utilize Principal Component Analyses to extract the most significant dynamical cross-correlations between these domains and explain them in terms of the glycan modifications. Finally, we end with a detailed analysis of epitope exposure and vulnerability at the conserved immuno-critical regions of the fusion peptide[44] and the heptad repeat regions[45, 46]. These results have equipped us with robust engineering capabilities to overcome vaccine and Spike-targeting therapeutic design challenges.

## RESULTS

The extensive MD simulations reported in this study include natively glycosylated Spike soluble ectodomain trimer of the G614 and Delta variants with the same glycoforms[39]. Each of these two variants was simulated in both all-RBD down (ACE2 inaccessible) and 1-RBD up (ACE2 binding compatible) conformations with five independent microsecond-long trajectories (total simulation time of 20μs; see SI Methods for details). In order to distinguish between the three similar protomers in the Spike, we will refer to them as chains A, B, and C (counter-clockwise arrangement), where chain B is open in the 1-RBD up system, chain A is to the left and chain C is to the right of the up chain B (see **Fig.1**).

### Enhanced Glycan Shielding of the NTD in the Delta Variant Reduces Antibody Accessibility

Here, we quantify the effective glycan shielding by employing the glycan-encounter-factor (GEF) metric designed by us earlier[30, 47], which essentially measures the relative glycan coverage encountered by an approach probe (an antibody, for example) over each residue on the protein surface, taking the glycan and protein dynamics into account. To provide an overview of how this glycan coverage may alter between variants, we calculated the residue-wise difference in GEF values (Delta minus G614) for the entire Spike (**Fig. S1**). The differences are most noteworthy in the NTD region, where glycan shielding across this domain is notably higher for the Delta variant, relative to G614 over a large area of contiguous protein surface (**Fig. S1**). To further illustrate this, **Fig. 2** shows the average GEF (calculated from all three protomers) as a function of NTD residues for the G614 and Delta simulations. In particular, NTD surfaces spanned by residues ∼100 to 190 and ∼230 to 280 exhibit significantly higher coverage by glycans in the Delta variant, as compared to G614 (**Fig. 2a/b/c**). We note that these NTD surfaces include residues from the NTD supersite and epitopes of most NTD-specific neutralizing antibodies[48, 49]. Hence, the increased glycan shielding in these NTD areas could help the Delta variant evade NTD-targeting antibodies.

**Figure 2.**
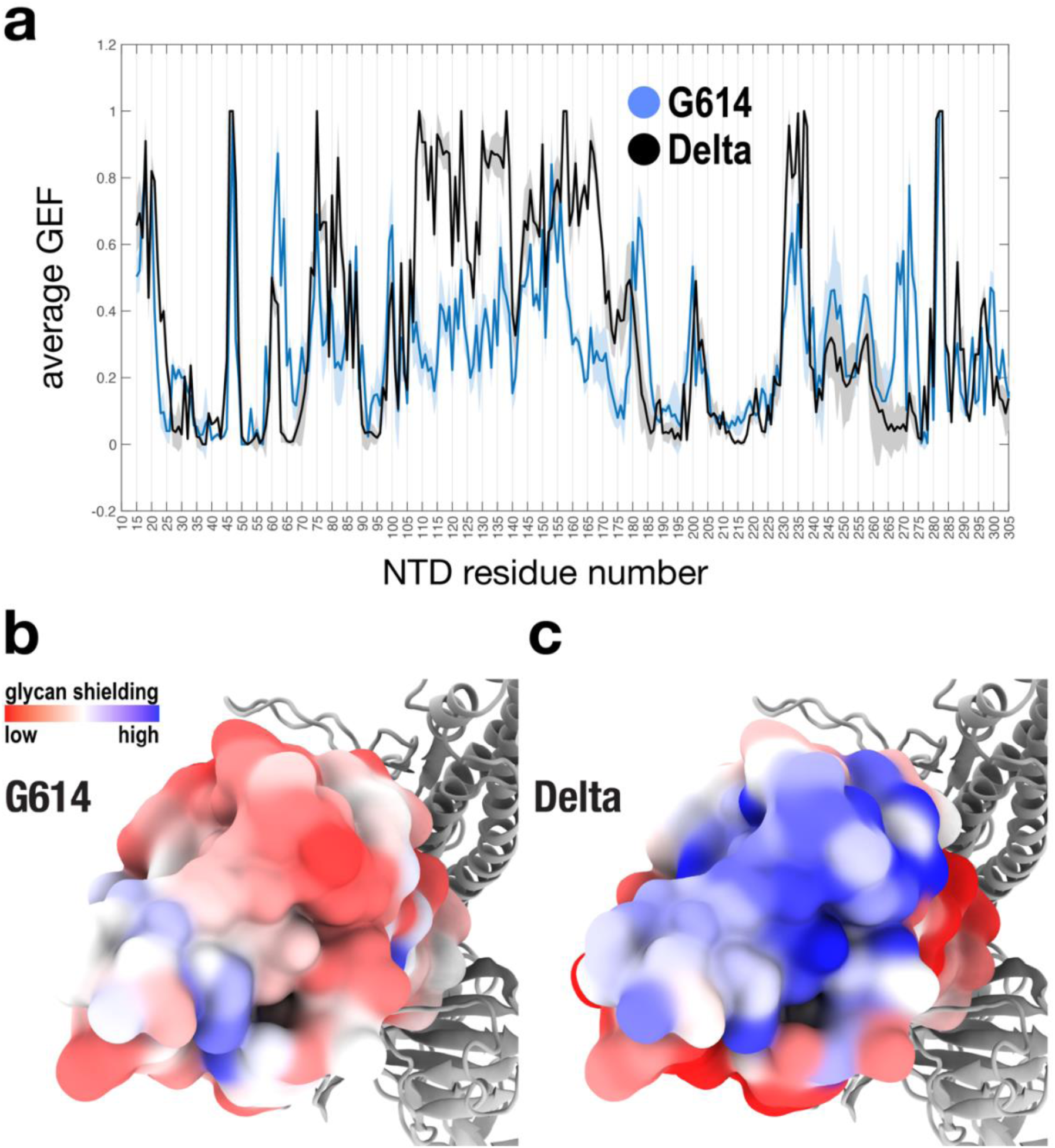
Glycan shielding of NTD surface. (**a**) To quantify glycan shielding, we calculated the glycan encounter factor (GEF) as function of NTD residue, for the G614 (blue) and Delta (black) simulations. The blue and black solid lines represent averages from all three protomers of the all-down G614 and all-down Delta systems, respectively. Filled error-bar envelopes show standard deviations. (**b/c**) To provide a structural interpretation, the average GEF values (as shown in panel **a**) are mapped onto NTD for (**b**) the G614 and (**c**) Delta variant. Qualitatively, GEF values are shown from low shielding (red) to high shielding(blue).

To elucidate additional contributors to NTD-antibody binding, we calculated the solvent-accessible surface area (SASA), for the NTD supersite that defines epitopes for most NTD-specific antibodies[49]. Changes in SASA would capture surface exposure variations because of protein structural burial. Interestingly, the SASA values remain similar in the NTD supersite for the G614 and Delta variants (**Fig. S2**). This suggests that the numerous modifications in the Delta NTD do not have any significant perturbation to the solvent-accessible surface area (SASA) of the NTD supersite. Rather, those Delta mutations modulate the orientations of the glycans, leading to enhanced glycan shielding of the NTD. Together, the results (cf. **Figs. S1, 2, S2**) corroborate the important role of the glycans that allow the Delta variant to become more evasive to NTD-directed antibodies. This finding is of critical concern since this corroborates and rationalizes previous evidence that NTD-specific monoclonal antibodies have reduced sensitivity for the Delta variant[50, 51].

### Restricted RBD Flexibility in Delta Facilitates ACE2 Accessibility

Since the viral RBD is responsible for binding to the host receptor, the dynamics of RBD are critical for how infectious a variant may be. To explore this aspect, we first checked the Root Mean Squared Fluctuations (RMSF) of the RBD residues (**Fig. S3**). Predictably, the overall RMSF of the up-RBD (protomer B) is significantly higher, in G614 (**Fig. S3b**). This is not unexpected since the RBD in the up conformation has a lesser number of interactions with neighboring protein domains to stabilize it. However, an interesting feature that emerges here indicates that the up-RBD is relatively more rigid (i.e. less flexible) in the Delta (**Fig. S3d**). To understand this difference further, we look into the essential dynamics of the RBD. We defined simple reaction coordinates that capture (i) an opening and (ii) a twisting movement of the RBD (see SI Methods and **Figs. S4, S5, 3**). To compare the simulated RBD dynamics, we calculated 2D distributions along RBD opening and twist, for all trajectories of G614 and Delta (**Figs. S4, S5, 3**). We find that the most significant difference between these variants is in the up-RBD conformation of protomer B (**Fig 3**). Specifically, the up-RBD of G614 samples a larger conformational space (leading to higher flexibility, **Figs. 3, S3**) than the up-RBD of Delta. Further, the dominant up-RBD ensembles (i.e. the locations of free-energy minima) differ between G614 and Delta, where the G614 up-RBD on average adopts greater opening and twisting angles than the Delta up-RBD, which is confined to a basin of smaller opening and twisting values (**Fig. 3**).

**Figure 3.**
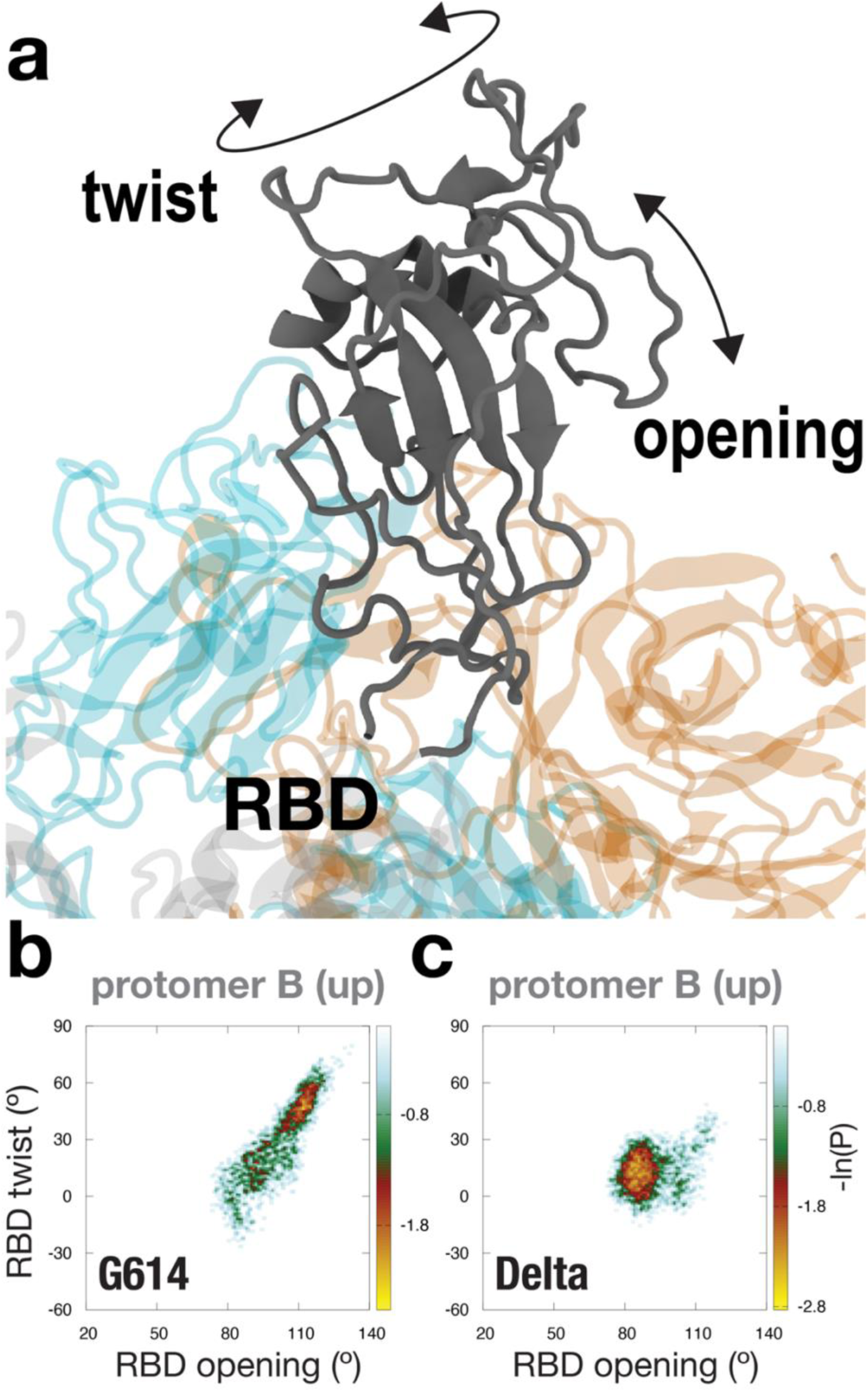
Movement of the up-RBD in protomer B, for 1-up G614 versus 1-up Delta. (**a**) To describe RBD rearrangements, we defined opening and twisting coordinates for RBD (see SI Methods for details). The up-RBD region is a zoomed-in view from Fig. 1a. (**b/c**) 2D distributions along RBD opening and RBD twist. The distributions in (**b**) and (**c**) describe the motions of the up-RBD in protomer B for the (**b**) 1-up G614 and (**c**) 1-up Delta systems, respectively.

To explore the biological implications of these RBD differences, we discuss how the distinct orientations of the up-RBD (in protomer B) may impact the opening movement of the neighboring RBD (in protomer A). For this, we chose two nearest-to-mean configurations where each represents the dominant up-RBD ensemble of the G614 and Delta simulations, respectively (**Fig. S6**). In these representative 1-up G614 and 1-up Delta structures, we then model an identical up-RBD in the neighboring protomer A, which is otherwise in the down orientation (see SI Methods). This allows us to infer how the up-transition of the protomer-A RBD may be affected by the variant-specific up-RBD orientations in the adjacent protomer B (cf. **Fig. 3**). In the G614 Spike, the up-RBD of protomer B, due to its the large opening angle and twist (cf. **Fig. 3b**), is positioned closer to the neighboring RBD of protomer A (**Fig. S6**). This apparently introduces steric hindrance that impedes the up-transition of the adjacent protomer-A RBD (**Fig. S6**). By contrast, in the Delta Spike, a smaller opening angle and twist (cf. **Fig. 3c**) allow the up-RBD of protomer B to remain further away from the neighboring RBD. This reduces potential steric clashes and likely facilitates the up-movement of the adjacent RBD in protomer A (**Fig. S6**). Together, these results expound the structural bases of observed Delta RBD structural diversity and increased population of 2-up or 3-up RBD configurations[52, 53], which would conceivably facilitate virus-host binding, thereby enhancing the infectivity of Delta, as compared to G614.

### NTD Tilting in Delta Influences RBD Orientation and Gating

Since each NTD can interact with an RBD of a neighboring protomer, it is instructive to also characterize the motion of NTD, in order to establish any correlated dynamics between adjacent NTD/RDB pairs. To this end, we defined structural coordinates that measure i) a tilting and ii) a torsion movement of the NTD (**Fig. 4a**). To compare the simulated NTD dynamics, 2D distributions along NTD tilting and torsion were computed for all trajectories (**Figs. S7, S8, 4b/c**). We find that the most notable difference between G614 and Delta is in the protomer C for the 1-up Spikes (**Fig. 4b/c**). The calculations demonstrate that the NTD can undergo large-scale rotational motions that are variant-specific. The G614 and Delta variants have distinct dominant NTD orientations in their respective protomer C (**Fig. 4b/c**) – the protomer where the NTD is structurally adjacent to the up-RBD (protomer B). Overall, the Delta NTD appears to sample a larger conformational space and adopts more tilted orientations, compared to G614 (**Fig. 4b/c**).

**Figure 4.**
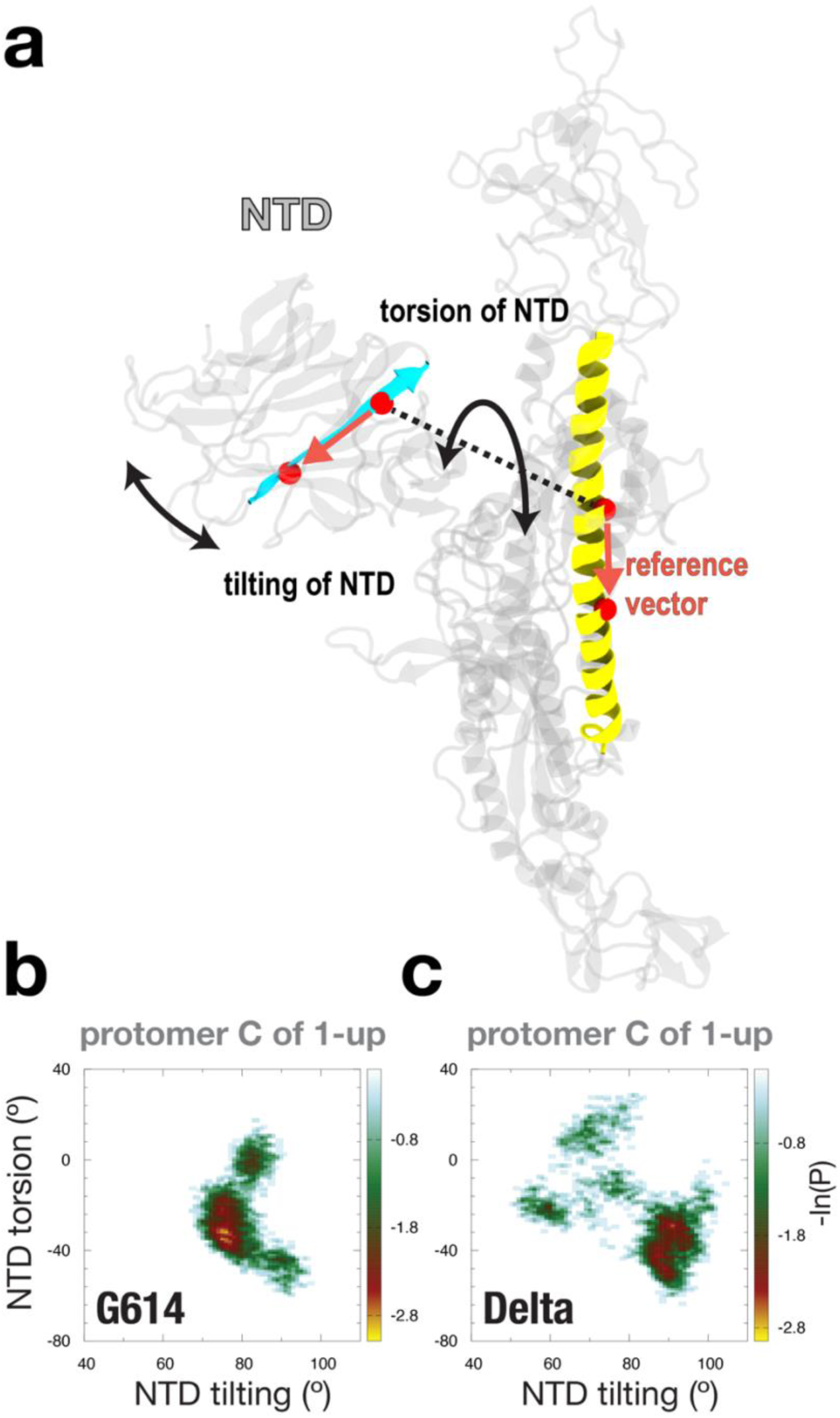
Movement of NTD in protomer C, for 1-up G614 versus 1-up Delta. (**a**) To describe NTD rearrangements, we defined tilting and torsion coordinates for NTD (see SI Methods for details). (**b/c**) 2D distributions along NTD tilting and NTD torsion. The distributions in (**b**) and (**c**) describe the motions of the NTD in protomer C for the (**b**) 1-up G614 and (**c)** 1-up Delta systems, respectively. Note that the NTD of protomer C is adjacent to the up-RBD of protomer B. Therefore, panels (**c**) and (**b**) are shown here to illustrate how the orientation of the protomer-C NTD may influence the orientation of the protomer-B RBD (cf. Fig. 3).

The variant-specific NTD dynamics in protomer C (of 1-up G614/Delta; cf. **Fig. 4b/c**) is related to the variant-specific RBD dynamics in protomer B (cf. **Fig. 3b/c**). To interpret this, note that the NTD of protomer C (or C-NTD) is nearest to the RBD up-protomer (protomer B or B-RBD). The other adjacent pairs are B-NTD/A-RBD and A-NTD/C-RBD (cf. **Fig. 1**), where A-RBD and C-RBD are RBD down-protomers in the 1-up Spike. Interestingly, the B-NTD and A-NTD, which are adjacent to RBD down-protomers, are overall more localized (i.e. less mobile), remaining at smaller NTD tilting and torsion angles (**Fig. S7**). This suggests that there is a stabilizing effect between an RBD down-protomer (**Fig. S4**) and its neighboring NTD (**Fig. S7**). By contrast, the C-NTD, adjacent to the up B-RBD, undergoes a wider range of tilt and torsion motions (**Fig. 4b/c**). Further, the distinct rearrangements of this C-NTD between G614 and Delta (**Fig. 4b/c**) appear to be associated with the variant-specific up B-RBD dynamics (**Fig. 3b/c**). This C-NTD/up B-RBD relationship is plausible because the numerous mutations in the NTD apparently alters the rearrangements of the NTD (cf. **Fig. 4b/c**), which likely impacts the orientation of its direct neighbor, the up-RBD (**Fig. 3b/c**).

### Glycan-Mediated Coupling of NTD and RBD Modulates Spike Conformations

Earlier, we discussed how the numerous modifications in the NTD can alter glycan orientations, leading to distinct glycan shielding of the NTD surface (cf. **Fig. 2**). In addition, the analysis of **Fig. 4** suggests that those NTD mutations also affect large-scale NTD rotations. These changes in NTD rearrangements (cf. **Fig. 4b/c**), together with the shift in glycan orientations (cf. **Fig. 2**), may influence the conformation of the neighboring up-RBD, which is characteristic of each Spike variant (cf. **Fig. 3**). To identify which glycans mediate the C-NTD/up B-RBD correlation (cf. **Figs. 3, 4**), we calculated the glycan conformational space for the dominant up-RBD ensembles (corresponding to the high occupancy basins) of the G614 and Delta trajectories (cf. **Fig. 3b/c**), respectively. We then visually compare the resultant glycan densities between those G614 and Delta ensembles in order to determine which glycans differ in orientation for different variants. Based on how they are positioned on the protein surface, these glycans may contribute to the correlated movement of the C-NTD and up B-RBD.

Near the up B-RBD, we find that glycans N165 and N234 in the adjacent C-NTD, and glycan N331 in the up B-RBD sample different orientations in the G614 and Delta simulations (**Fig. 5**). Specifcally, a comparison of these simulations suggests that these three glycans cling to (and move with) the up B-RBD, which appears to accommodate (and influence) the specific up B-RBD orientations observed in the G614 and Delta simulations (**Fig. 5**). Glycans N165 and N234 have been previously shown to strongly affect Spike opening[15], with N165 increasing and N234 decreasing RBD up conformations[54] in the original Wuhan strain. Even glycan N331, located at the base of the RBD, can affect receptor binding[55] and antibody response[56]. Our simulations indicate that these glycans lodge at the base of the RBD in different orientations, and stabilize disparate RBD conformations.

**Figure 5.**
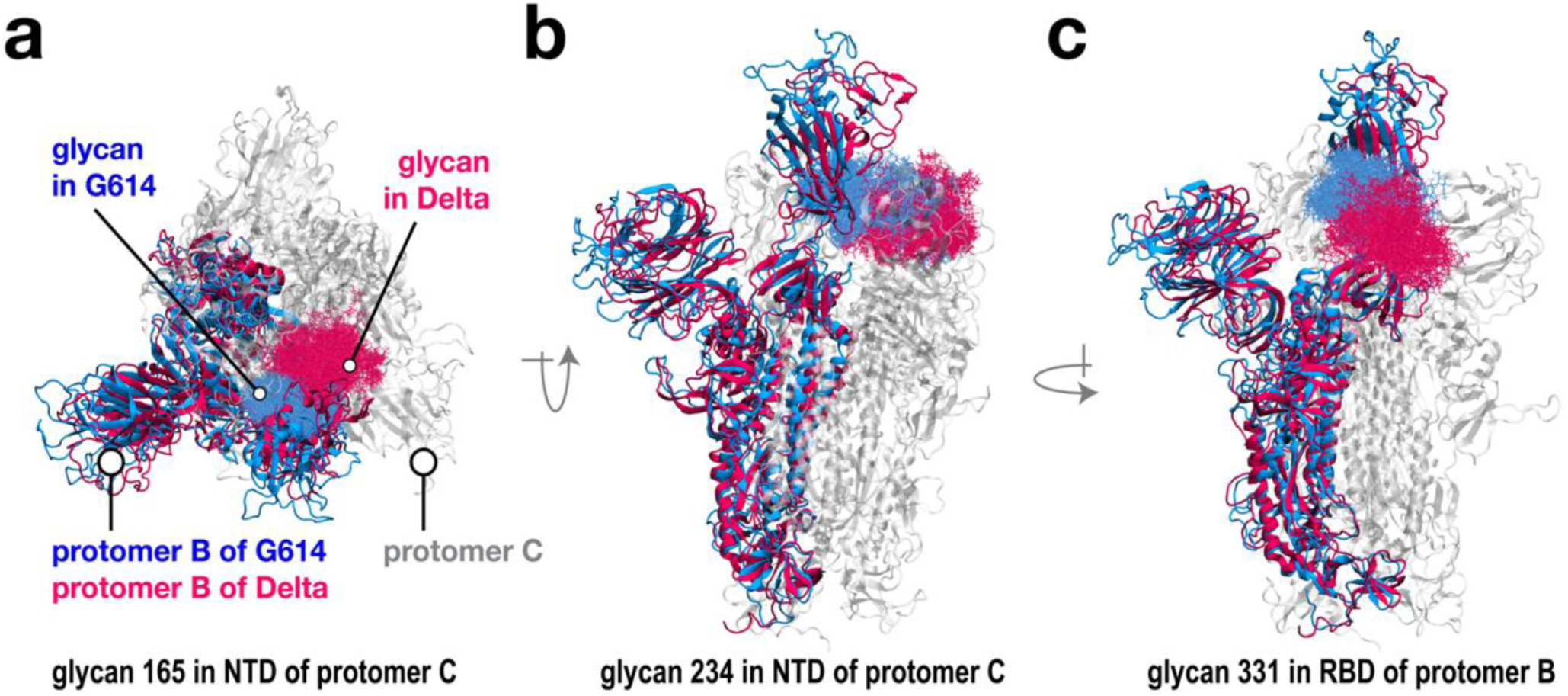
Distinct glycan positioning in G614 versus Delta may lead to distinct up-RBD orientations. Near the up-RBD of protomer B, the simulations suggest that glycans (**a**) 165 and (**b**) 234 in the NTD of the neighboring protomer C, and (**c**) glycan 331 in the RBD of protomer B adopt distinct orientations in the G614 (blue) versus Delta (magenta) simulations. In all panels, the overlayed blue and magenta structures are representative conformations of protomer B from the dominant up-RBD basins of the G614 (i.e. blue) and Delta (i.e. magenta) simulations, respectively (cf. Fig. 3b**/c**). In each panel, the neighboring protomer C is shown in orange, for reference. In panels **a/b/c**, blue point densities depict the positions of glycans (**a**) 165, (**b**) 234, and (**c**) 331 in the G614 simulations. Similarly, magenta point densities depict the positions of glycans (**a**) 165, (**b**) 234, and (**c**) 331 in the Delta simulations.

These results demonstrate how mutations in the NTD, and glycans, can have far-reaching implications for the function of the Spike. That is, changes in NTD rearrangements can not only directly affect the RBD motion of neighboring protomers, but also influence the glycans at the interface of these two critical domains. These differentially oriented glycans can aid in maintaining altered RBD conformations between variants.

### RBD-gating motion is allosterically modulated by the adjacent NTD

There has been a putative conjecture that the structure and dynamics of the NTD may be allosterically connected to RBD gating[40, 57]. Given the distinctive RBD and NTD rotational conformations observed above and the notable reorientation of key glycans between these two critical domains (cf. **Figs. 3, 4, 5**), we set out to study the cross-correlated fluctuations that may drive large-scale allosteric motions within the peplomer. The inter-residue dynamic cross-correlation plots (**Figs. S9, S10**) indicate that beyond the expected strong correlations within individual NTD and RBD domains, there is a remarkable inter-domain NTD-RBD synchronization of dynamics as well. One must note that in the closed conformation, the RBD of one protomer directly interacts with the NTD of the protomer situated anti-clockwise to it (cf. **Fig. 1a**). In our findings, as seen in **Figs. S9a/b,** the RBD of chain A is correlated with the NTD of chain B, the RBD of chain B is correlated with the NTD of chain C, and the RBD of chain C is correlated with the NTD of chain A in both G614 and Delta closed conformations. More remarkably, the up-RBD (chain B), even in the open conformation, has strongly correlated fluctuations with the neighboring NTD (chain C), which is much stronger in the Delta variant as compared to G614 (**Fig. 6a/b**). Once the RBD moves up, it is no longer in direct interaction with any neighboring domain. Yet this strong correlation of motion suggests allosteric communication between the up-RBD and the NTD to its right. Recollecting the role of glycans N165 and N234 from that NTD and N331 from the base of the up-RBD in propping the receptor binding domain in distinct conformations (Fig. 5), these glycans may be the key to NTD-RBD cross-talk over and above communications via the protein backbone. This strong correlated motion between inter-protomer NTD and RBD in the up-conformation and its relative increase in strength in Delta establishes the critical role of NTD in RBD gating and indicates that this allosteric communication is better optimized in the Delta variant.

**Figure 6:**
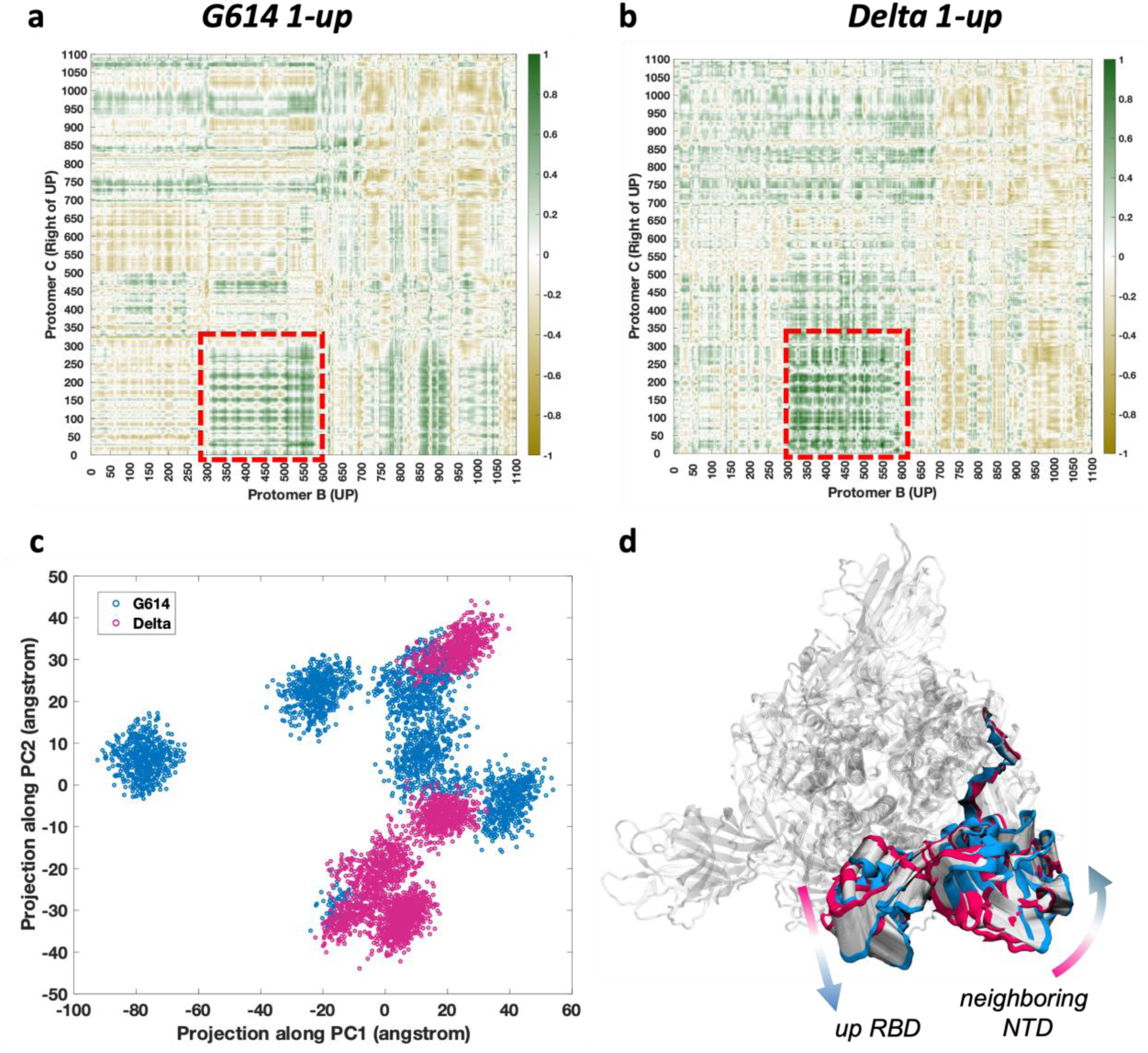
Differences in correlated motion of up-RBD with neighboring NTD, between G614 and Delta. (**a**) Inter-residue cross-correlation of RMS fluctuations between protomer B (RBD-up) and Protomer C (right neighbor) in G614 (1-up), and (**b**) Delta (1-up) conformations. Interprotomer RBD-NTD correlation space is indicated by red box. (**c**) Projection of the G614 (blue) and Delta (magenta) ensembles along the top two principal component eigenvectors. PC analysis was performed over the two combined ensembles to extract the variances in large-amplitude motions (see methods for details). (**d**) Structural representation of motion along PC2, which describes the dominant difference in motion between G614 and Delta. Magenta structure represents negative PC2 (Delta-like) and blue represents positive PC2 (G614-like), with interpolation given in white. Motion of the up-RBD (protomer B) and neighboring NTD (protomer C) is shown here.

To extract the large-amplitude low-frequency dynamics that result from these correlated fluctuations, we performed a Principal Component Analysis (PCA) over the simulated ensemble (see details in SI Methods). We extracted the top two PCs that maximize the variance in motions and that define the majority of the essential dynamics that drive Spike equilibrium motions. These PCs in both up and down conformations indicate that in equilibrium conditions, the dominant motions are mainly confined at the bulk-head of the Spike, within the NTD-RBD regions – at least within the microsecond time-scale sampled in our simulations (**Fig. S11**). Let us independently consider the down and up configurations of the Spike. With all RBD down and closed, there is a distinct separation in RBD-NTD orientation between G614 and Delta variants (Fig. **S9 c/d**) along PC1. The RBD is rotated closer to the central axis in Delta, with a concomitant rotation of the neighboring NTD inward towards the RBD. The PC2 motion remains invariant between G614 and Delta, indicating that NTD-RBD correlated allostery has a variant-independent component. Looking at the up-conformations next (**Fig. 6c**), we find a similar variant independent NTD-RBD allosteric component, which forms the essential motion with the highest contribution (i.e., along PC1). PC2, however, shows a significantly high occurrence of separation between G614 and Delta, where the up-RBD is tilted more inwards towards the central axis in Delta, accompanied by the NTD tilted downward towards the stem of the Spike (**Fig. 6d**). This correlated motion is similar to the combination of RBD and NTD orientational variations discussed above (cf. **Figs. 3, 4**), establishing that the RBD opening-twisting motion and the NTD tilting motion are allosterically coupled, further implicating the role of NTD in Spike gating.

### D950N Mutation in Delta Alters HR1 Dynamics and Spike Stability

Beyond the RBD mutations that are expected to have direct ACE2 binding and antibody escape effects[37], the FCS mutation with furin cleavage effect[58], and the other varied NTD/RBD mutation effects as discussed above, the Delta variant also has the interesting D950N substitution. This is structurally located in the S1-S2 interface and at the highly conserved heptad repeat (HR1) (cf. Fig. 1). To shed light on the evolutionary advantage of having this mutation on an otherwise conserved helix, we performed a detailed structural analysis of this region. To elucidate the impact of D950N, it is instructive to identify any residues that interact with the mutation site 950. For this, we calculated the probability of contacting residue 950 for all other residues in the Spike, using the G614 and Delta simulations (**Fig. S12**). We find only two residues (i.e. E309 and R1014; see **Fig. S12**) that form noticeable contacts with 950, in both the G614 and Delta systems. Because these residues are negatively (E309) and positively (R1014) charged, their interactions with 950 are expected to alter since the D950N mutation removes the negative charge. To discern possible changes in these interactions, we calculated distances between (i) 950 and 309, and (ii) 950 and 1014, for G615 (with D950) versus Delta (with D950N). The results indicate that the D950N mutation leads to distinct distance distributions, especially when comparing the up-protomers of G614 against Delta (**Figs. S13, S14**). These changes suggest that the HR1 may undergo variant-specific fluctuations.

To properly capture the dynamics of HR1, we note that the HR1 consists of two helices as shown in **Fig. 7a and 7b**, which are in the regions ∼910-941 (referred to as “HR1-bottom”) and ∼942-986 (“HR1-top”). Since HR1-bottom and HR1-top may undergo independent rearrangements from each other, we define separate sets of coordinates that describe the movement of each part individually. That is, for each HR1 part, we introduced a tilting and a torsion coordinate, which are measured relative to the central helix (CH) of the Spike (see SI for details; **Figs. S15-S18**). For the HR1-top region (which has the mutation site D950N), the most noticeable difference between G614 and Delta is in the fluctuation along the torsion coordinate, in the up-protomer (**Fig. S15, S16**). Specifically, HR1-top of G614 exhibits a larger range of torsion angles in the up-protomers, while the torsion range in the remaining down-protomers is smaller. In contrast, the torsion range of the HR1-top in Delta remains smaller in all protomers, regardless of up or down (**Fig. S15, S16**). For the HR1-bottom region, the most dominant difference between G614 and Delta is in the rearrangement along the tilting coordinate, in the up-protomer (**Fig. S17**). Specifically, the HR1-bottom of the up-protomer in Delta can reach significantly larger tilting angles (max tilt ∼30°), compared to the G614 simulations (maximum tilt ∼20°).

**Figure 7.**
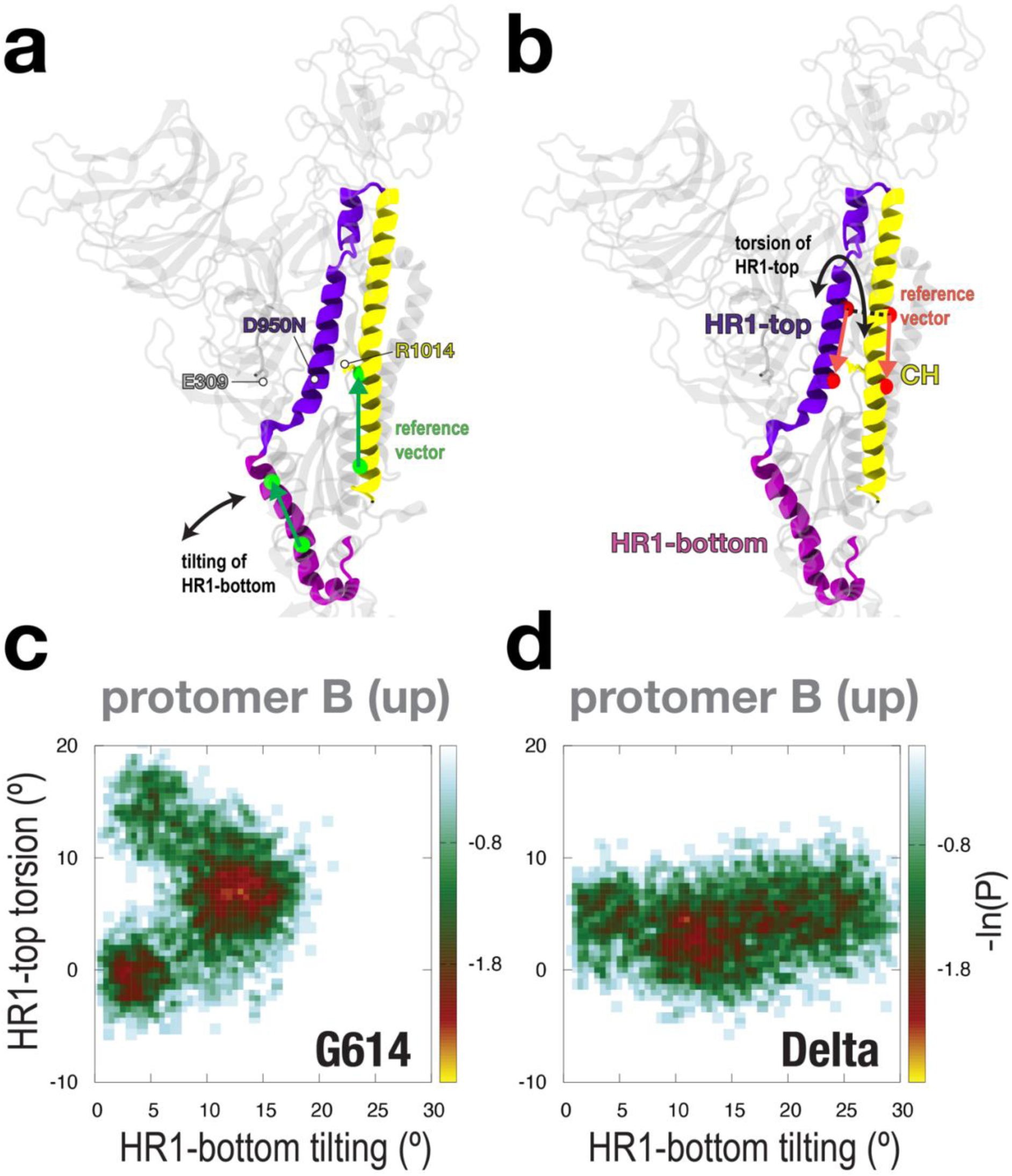
Movement of HR1 in protomer B, for 1-up G614 versus 1-up Delta. (**a/b**) To describe HR1 rearrangements, we capture motions in the bottom (purple) and top (violet) part of HR1. Specifically, we measure (a) a tilt of HR1-bottom relative to CH (yellow) and (b) a torsion of HR1-top relative to CH. See **Figs. S15** and **S17** for details on how these coordinates are defined. (**c/d**) 2D distributions along HR1-bottom tilting and HR1-top torsion. The distributions in (**c**) and (**d**) describe the motions of the HR1 in protomer B for the (**c**) 1-up G614 and (**d**) 1-up Delta systems, respectively.

Together, the analyses of **Figs. S15-S18** allow us to identify the relevant motions of each HR1 part, along which changes between the variants occur. To summarize these changes, we plot 2D distributions along (a) the torsion of the HR1-top and (b) the tilt of the HR1-bottom, which encapsulate the key differences between the G614 and Delta simulations (**Fig. 7**). In conclusion, the results demonstrate that, specifically for an up-protomer, the D950N mutation can be responsible for complex changes in HR1 rearrangements (**Fig. 7**), which may influence how glycans shield this region as discussed below (cf. **Fig. 8**), or even influence Spike-host fusion events.

**Figure 8.**
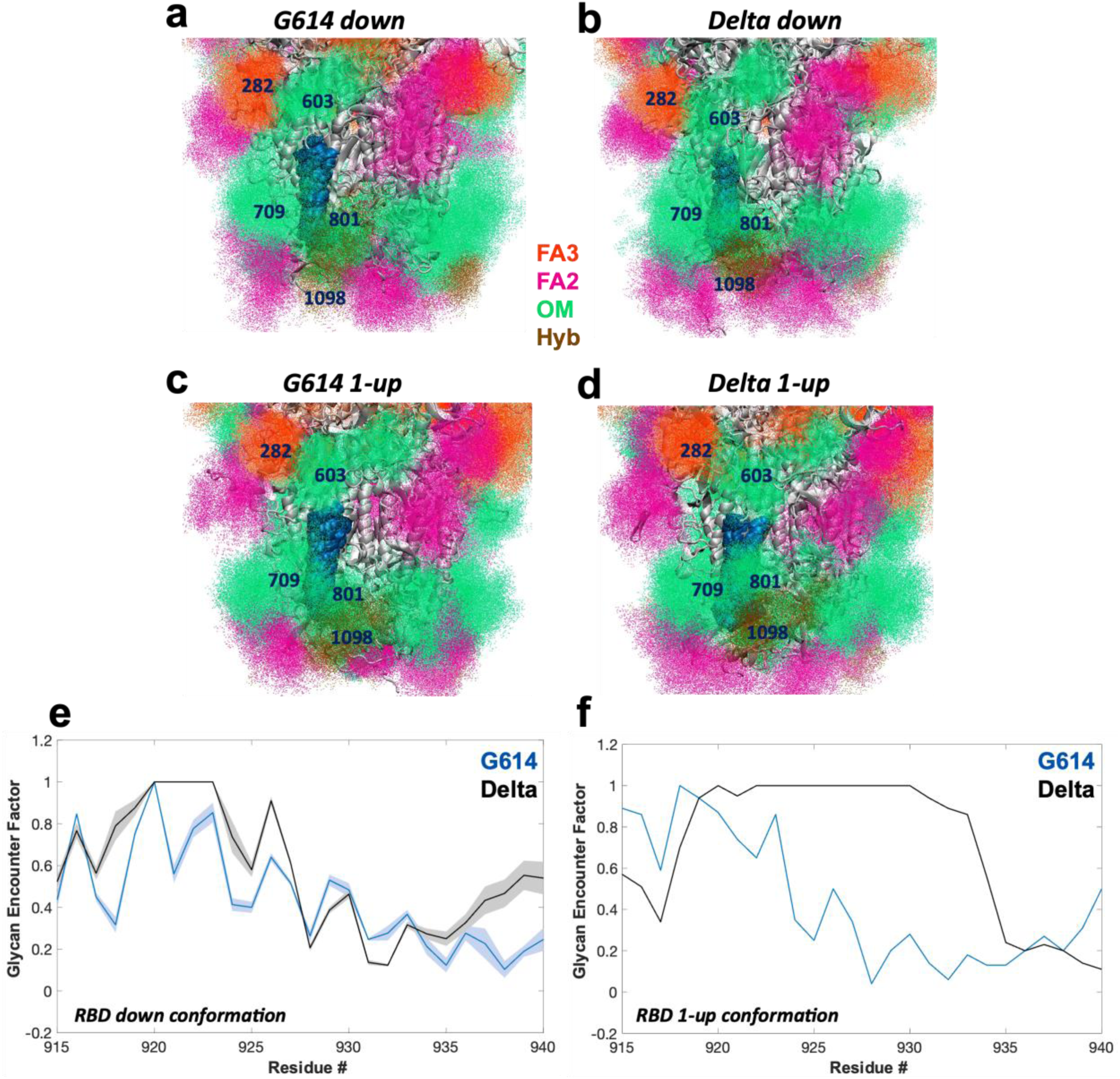
Glycan shielding of the HR1 epitope (residues 915-940) region. (**a**) G614 all-down, (**b**) Delta all-down, (**c**) G614 1-up, and (**d**) Delta 1-up conformations of the fusion peptide region. Multiple conformations of the fusion-peptide residues 800 to 860 are shown in dark blue, and the rest of the protein is shown in gray. Glycans near this region are represented by density of points, and colored according to the labeling. (**e**) Residue-wise GEF plot for G614 (blue) and Delta (black) Spikes in the RBD down conformation. Mean GEF values from the three protomers are shown, with standard deviation. (**f**) Residue-wise GEF plot for G614 (blue) and Delta (black) Spikes in the RBD 1-up conformation. GEF values only from the up protomer are shown.

### Glycan Shielding of the Fusion Peptide and HR1 is Enhanced in Delta

The fusion peptide (FP) region consists of sequences (800-860) that are highly conserved and are a hotbed of broadly neutralizing antibody targets and vaccine-design efforts[44, 59]. While the sequences in the FP region are conserved, mutations from other parts of the Spike can alter the shape, fluctuation, or glycan shielding of the FP region. A detailed understanding of these aspects would enable future experiments to develop more precise therapeutics.

Since glycan shielding is a critical factor for antibody evasion, we have identified which glycans are responsible for shielding the FP region. We have also examined if and how the glycan shielding may change (i) for different variants (i.e. G614 vs. Delta) and (ii) for different protomer conformations (i.e. all-down vs. 1-up). For the 800-860 region, we calculated the glycan coverage for the all-down G614, all-down Delta, 1-up G614, and 1-up Delta simulations (**Fig. S19a-d**). For all four simulation systems, glycans 603, 283, and 709 are the dominant contributors to shielding the FP. To quantify the extent of glycan shielding, we calculated the glycan-encounter-factor (GEF)[47] for each residue of this region (**Fig. S19e/f**). Overall, residues 805-825 receive on average less glycan shielding, while 800-803 and 830-855 receive greater shielding. Interestingly, this GEF profile remains similar across different protomer conformations, as well as variants (**Fig. S19e/f**). Hence, these findings suggest that glycan shielding of the FP region is a robust feature of Spike function, with glycans 603, 283, and 709 being the key contributors.

Given the cross-reactive human monoclonal antibody response found to be mounted over different sarbecovirus Spikes at the HR1 domain[45, 46], and the Delta-specific D950N mutation and causing dynamic variations in this neighborhood, we also performed a similar glycan coverage analysis for the 915-940 region (**Fig. 8**). All four simulation systems (i.e. all-down G614/Delta and 1-up G614/Delta) indicate that glycans 603, 801, and 1098 are the dominant contributors to shielding the 915-940 region (**Fig. 8a-d**). As for quantifying the extent of glycan shielding, the GEF profile is overall similar between the all-down G614 and all-down Delta simulations (**Fig. 8e**). Specifically, in each down-protomer of either variant (**Fig. 8e**), residues 915-925 receive overall greater glycan shielding, while residues 930-940 exhibited lower glycan shielding. Interestingly, this GEF profile remains overall similar in the up-protomer of the 1-up G614 simulations (**Fig. 8f**). However, the GEF profile changes significantly in the up-protomer of the 1-up Delta simulations. Specifically, the glycan coverage around residues 925-935 is noticeably higher in the up-protomer of Delta, as compared to the G614 variant (**Fig. 8f**). In summary, the glycan shielding of the 915-940 region in the G614 variant is independent of protomer conformation (cf. **Fig. 8e/f**). By contrast, glycan shielding increases for the Delta variant if the protomer is in the up-RBD conformation (cf. **Fig. 8e/f**). In particular, this increased shielding is attributed to the contributions from glycans 801 and 1098 in the up-protomer state, while the contribution from glycan 603 remains effectively constant between protomer conformations.

## DISCUSSION

Being exposed on the surface of the virus and forming the first line of contact with the host, different modalities of vaccine design, immune-mediated therapeutics, and even many small-molecule drugs primarily revolve around the SARS Spike glycoprotein. A molecular-level understanding of the mechanism of immune evasion, as well as the dynamic cascade of events that lead to functional interaction with the ACE2 receptor, is needed to get a better understanding of this critical protein. However, due to structural complexities, as well as constantly evolving antigenic surfaces, the structure-dynamic relationships of this glycoprotein are not easily attainable through experimental studies. As utilized here, a computational approach provides strong alternatives to augment experimental findings. Even though the SARS-CoV-2 virus has shown an impressive number of variants accumulating over time from its emergence in late 2019, the D614G substitution was one early natural selection[29] that has stably persisted through subsequent evolutionary events. On the other hand, the Delta variant emerged as one of the most virulent, with high symptom severity among the VOCs[9]. The Delta variant serves as an ideal model for studying the interplay between infectivity, immune evasion, and structural adaptability. Many of the key mutational effects observed in Delta remain relevant, as similar alterations persist in Omicron-derived sublineages[60] and later variants. Additionally, immune escape mechanisms first observed in Delta shaped the trajectory of SARS-CoV-2 evolution, reinforcing its role as a valuable reference for future studies on viral adaptation and immunogen design[61]. Here, to illustrate allosteric coupling mechanisms and antibody evasion strategies that provide fitness advantages to a specific variant, we perform a comparative examination of the D614G and Delta (B.1.617.2) Spike dynamics , as well as quantify the intrinsic Spike properties that have remained invariant despite evolution.

Of the ∼22 glycans present on an individual Spike protomer, nine are present on the NTD, one on the RBD, and one on the connector between these two domains. Like most peplomeric glycans, one of their primary purposes is shielding the underlying antigenic surface from antibody interaction. The Spike NTD houses the epitopes of several broadly neutralizing antibodies, instigating a repertoire of mutations and deletions, many of them within the residue ranges of 14-26, 141-156, and 246-260, commonly known as the NTD supersite due to their high antibody sensitivities[49]. This study demonstrates that the NTD glycans, specifically N17, N149, N165, N234, and N282, are reorganized in their spatial orientation, possibly through the 5 Delta mutations and two deletions in this domain (see **Fig. 1b**). Consequently, the shielding effect increases over the Delta NTD, especially around the supersite residues (**Fig. 2, S1**). This explains the reduced neutralization capability of NTD-specific antibodies to the Delta[50, 51], allowing this variant to evolve into a more immune-evasive variety. While most studies target the direct local interactions of the substituted amino acids with antibodies to explain changing immune responses, this finding shows that it is essential to take the local glycan orientation into account as well.

The upward gating of the RBD is an essential dynamic transition that causes ACE2 binding and subsequent fusion events. However, the RBD-open conformation also allows for antibody targeting that competes with the host receptor. Our findings establish that the Delta RBD, on average, adopts a less erect poise, having lower ‘opening’ and ‘twisting’ angles as compared to the G614 RBD (**Fig. 3**). From our analysis, it is evident that this structural basin concedes greater steric clearance from neighboring domains (**Fig. S6**), allowing for multiple RBDs to open simultaneously, which has already been evidenced in experiments[62]. This greater separation between neighboring up-RBDs also justifies the hitherto unexplained multiple ACE-2 associations observed in Delta without expected steric clashes[24]. We speculate that this differently poised up-RBD probability may also aid in avoiding receptor binding motif targeting antibodies while allowing for multi-valent ACE2 interactions.

Complementing the differential stance of up-RBD, our results also illuminate a differently oriented NTD with a higher probability of being more tilted towards the Spike stem (**Fig. 4**) in Delta. We speculate that the multiple amino acid substitutions and deletions alter the local structural arrangements in the NTD, which in turn affects the dynamics of this domain, making it take up an altered tilt, as well as reorient the NTD glycans and consequent shielding, as discussed before. The exact role of the large globular NTD in the Spike function has been elusive. Many antibodies have been found to target the NTD, rather than the ACE2-binding motif, yet it is unclear how binding to this distant domain results in Spike neutralization[13]. This indicates that preventing the normal dynamics of the NTD with the bulky attachment of antibodies somehow arrests the downstream functioning of the Spike. Keeping in mind that the RBD of one protomer is in physical interaction with the neighboring protomer NTD (situated anticlockwise, to the right), our conjecture is that the tilting and torsional motion sampled by the NTD dynamics (**Fig. 4**) triggers RBD opening in the directly neighboring protomer.

Another noteworthy finding has been the effects of a trifecta of glycans around the open RBD (in protomer B). Glycans N165 and N234 from the adjacent NTD (in protomer C) and glycan N331 from the base of the up-RBD have altered orientations in terms of overall sampled space (**Fig. 5**). It has been previously understood that beyond shielding, glycans also aid with RBD transitions[42]. Here, we further demonstrate that the glycans also play vital roles in locking the RBD in specific positions by nudging against the RBD base. These interfacial glycans not only help with optimized glycan shielding but also influence the RBD conformations by helping the domain maintain varied structural stances – in the case of Delta, allowing for a more bent conformation towards the central Spike axis.

Given the NTD and RBD dynamics sample altered landscapes for G614 and Delta Spikes, and that glycans can reorient and affect domain dynamics between these variants, we have also elucidated the correlated motions and large-amplitude essential dynamics captured in the simulations. Our findings validate that the NTD and RBD demonstrate the largest motions along the high-contribution variational coordinates. Moreover, beyond the intra-domain correlations, the inter-protomer NTD and RBD show strongly correlated motions. Most noteworthy is the coupling between the up-RBD and the neighboring NTD (anti-clockwise) that becomes stronger for Delta. This validates our conjecture that the motions of the NTD may trigger the adjacent RBD dynamics. PCA demonstrates that both closed and open Spikes tend to occupy separate RBD-NTD relative orientation and rotational conformations between the two variants (**Fig. 6, S9**). These different orientations, also captured by our reaction coordinate analyses (cf. **Fig. 3, 4**), are strongly coupled, as evidenced by the high eigenvalues of the PCA. Yet, one of the top two eigenvalue components (PC1 for down-RBD and PC2 for up-RBD) is variant-independent. These motions, therefore, extract the invariant allosteric signaling between NTD and RBD that is intrinsic to the Spike functioning – demonstrating the dynamic role of NTD in the RBD gating action.

Similar mutation-induced allosteric alterations are also possible in other regions of Spike as highlighted by another vital mutation of the Delta that is interestingly situated away from the RBD or NTD, which is the D950N substitution on the otherwise conserved HR1 helix. Our simulations demonstrate the D to N substitution in this position, which removes a negative charge, affects the local electrostatic interactions with E309 and R1014. As a result, the HR1 dynamics is altered significantly (**Fig. 7**), with the top half of the HR1 region having greater torsional motion in G614 and the bottom half having a greater tilting motion in Delta. These dynamic changes affected the exposure of the HR1 epitope region, where the glycan shielding is increased in the Delta RBD-up conformation. Located at the interface of S1/S2, these dynamic variations can also influence fusion with the host membrane, possibly enhancing the fusion ability in Delta[62].

In summary, this study has elucidated the allosteric modulation of Spike RBD by the N-terminal domain of the neighboring protomer. This dynamic coupling has both a variant-dependent and an invariant component. Delta-specific mutations on the NTD cause local structural changes that affect the rotation and tilting of the NTD. This has two important consequences critical to Spike’s behavior. The glycans on the NTD get differently oriented and cause greater antigenic surface shielding, rendering the Delta variant less sensitive to NTD-specific antibody neutralization. Secondly, this altered rotational dynamics of the NTD is allosterically communicated to the neighboring RBD, causing it to, in turn, sample a modified twist and opening conformation. A trio of glycans engages the RBD in this changed conformation, which allows for reduced steric clash and multiple RBD-open modes. A cryo-EM study has demonstrated that the Delta Spike exhibits extensive diversity in open-close substates compared to other variants [52] Besides the mutation-dependent RBD-NTD communications, there are variant-independent allosteric dynamics that lead the NTD bulk-head motions to trigger the RBD-gating mechanism under equilibrium conditions, without external forces. The D950N mutation at the HR1 further affects the S2 helix dynamics, governing the glycan coverage in the HR1 epitope region.

The antigenic nature of the Spike, along with its critical functional aspects of receptor binding, as well as the plasticity of multiple conformations, make this protein an attractive target for drug and vaccine development studies targeting potential human coronaviruses. When properly understood, its functional versatility and complexity can be harnessed to understand evolutionary strategies of other coronaviruses. With control over variant-specific structural coupling (e.g. differential NTD-RBD relative orientations) and glycan coverage (e.g. over NTD supersite and the conserved HR1) as well as the invariant allostery (e.g. NTD-triggered RBD gating) and epitope shielding (e.g. over the flexible fusion peptide region), both VOC-specific as well as generalized sarbecovirus immunogen strategies can be engineered. Targeting allosteric pathways that regulate RBD gating—rather than just the ACE2-binding interface—could be a promising avenue for designing broadly effective inhibitors. Additionally, understanding glycan shielding at the NTD and HR1 regions provides a rationale for optimizing monoclonal antibody cocktails that account for glycan-induced epitope occlusion.

## Funding and Acknowledgements

SC was partially supported by Los Alamos National Laboratory (LANL) Center of Nonlinear Studies Postdoctoral program. KN and SG were supported by NIH/NIAID/DMID grant P01 AI158571. This research used computational resources from LANL Institutional Computing Program and Northeastern Discovery cluster at MGHPCC. SC would also like to acknowledge the NEU Startup Faculty Funds. The authors acknowledge and thank Bette Korber, LANL, for her invaluable insights about HIV Env immunogenic structure and function.

## Author contributions

S.C., K.N., and S.G. conceptualized the study and designed the experiments. S.C. and K.N. prepared the structures. SC ran the simulations. S.C., K.N., and S.G. performed the data analysis and interpretation. S.C. and K.N. prepared the figures. S.C., K.N., M.Z, and S.G. wrote and edited the manuscript. S.C. and S.G. are co-corresponding authors.

## Data and materials availability

Data and materials are available from the corresponding authors upon request.

