## Supplementary material for "Allosteric Control and Glycan Shielding Adaptations in the SARS-CoV-2 Spike from Early to Peak Virulence": Methods and Supplemental Information

### METHOD DETAILS

**Model setup and visualization:** An optimized soluble G614 Spike structure as extensively tested by a previous study in the group was used as the starting conformation[1]. In short, a structural threading from Wuhan strain cryo-EM structure[2], along with other SARS-CoV1 based templates for missing regions was performed by homology modeling. The model was further improved by flexible fitting into cryo-EM maps[3], followed by large scale MD simulation runs. The closest to mean structure for G614 variant was selected for this study. Delta mutations were added by homology modelling in MODELLER 9.10[4]. Site-specific glycoforms were selected from the most probable distributions as determined in mass spectrometry studies[5], and were kept the same between the D614 and Delta variants. Glycan addition and relaxation was performed using the ALLOSMOD protocol[6]. All visualizations and structural representations were performed with VMD 1.9.3[7].

**Simulation Protocol:** All-atom explicit-solvent simulations were performed with the AMBER-16 software package[8, 9]. The CHARMM36m protein[10] and CHARMM36 carbohydrate[11] forcefields were used, with the TIP3P water model[12]. Each Spike configuration was centered in a cubic box and hydrated. The size of the box was chosen to create at least 15 Å padding on each side along the largest atom-atom distance of the Spike-glycan complex. The system was neutralized with an excess of 150 mM KCl. Each simulation was energy minimized using steepest descent and equilibrated in three stages. The first equilibration stage involved restrained simulation in the constant number-volume-temperature (NVT) ensemble for 2 ns. The second stage involved restrained simulation in the constant number-pressure-temperature (NPT) ensemble for 10 ns. During the first and second stages, harmonic position restraints were imposed on all non-hydrogen atoms. The third stage of equilibration involved 10 ns of simulation in the NPT ensemble and only included position restraints on protein backbone atoms.

A constant temperature of 310 K was maintained using velocity Langevin dynamics[13], with a relaxation time of 1 ps. A constant pressure of 1 bar was maintained using the Berendsen barostat[14], with a relaxation time of 4 ps and compressibility of $4.5 \times 10^{-5}$ bar$^{-1}$. Covalent bond lengths involving hydrogens were constrained using the SHAKE algorithm[15], with a tolerance of $10^{-6}$ nm. Water molecules were rigidified with SETTLE[16]. Lennard Jones interactions were evaluated using a cutoff where forces smoothly decay to zero between 1.0 and 1.2 nm. Coulomb interactions were calculated using the particle-mesh Ewald (PME) method[16], with a Fourier grid spacing of 0.10 nm and fourth order interpolation. Unrestrained production simulations were performed in the NPT ensemble, with an integration timestep of 4 fs that was enabled through hydrogen mass repartitioning[17].

Simulations were run for four systems: 1) G614 all-down, 2) G614 1-up, 3) Delta all-down, and 4) Delta 1-up. For each system, 5 replicas were generated, where each replica is 1.2 $\mu$s long. The first 200ns was the equilibration phase, and the final 1 $\mu$s was used as production ensemble. Accordingly, each system has an accumulated length of 5 $\mu$s of production ensemble. In total, 20 simulations were performed for an aggregated time of 20 $\mu$s. The G-form simulations have also been utilized in another independent study[18].

**Glycan Encounter Factor (GEF):** Glycan shielding effect over the protein surface was calculated per-residue, based on the glycan encounter factor (GEF) score[19]. This was calculated as the geometric mean of the probability that a probe approaching a surface residue would encounter glycan heavy atoms, in perpendicular and tangential directions. Probe size of 6 angstrom diameter was chosen to mimic a typical hairpin loop. Calculations were performed in x-y-z cardinal directions and their geometric mean was taken and normalized. Computational modeling calculations were implemented with VMD 1.9.3[7], Python[20] and MATLAB 2018[21].

**Root mean square fluctuation (RMSF):** Root mean square fluctuation (RMSF) of atom *i* is calculated as,

$$RMSF_i = \sqrt{\frac{1}{T}\sum_{t_j=1}^{T}\left\|\boldsymbol{r}_{\mathbf{i}}(\mathrm{t_j}) - \boldsymbol{r}_i^{\mathbf{ref}}\right\|^2}\ ,$$

where $T$ is the total time of the production run, tj is the index for time point of calculation, $\boldsymbol{r}_i^{ref}$ is the coordinates of the mean structure, and $\boldsymbol{r}_i$ is the coordinates of the *i*th frame. All calculations were performed with the biopython libraries[22].

**Coordinates for RBD rearrangements:** To describe RBD movement, we define an opening and a twist motion of the RBD (cf. **Fig. 3**), which are measured relative to the central helix (CH) (see

**Fig. 7b**) and upstream helix (UH). To this end, we introduce a reference vector (spanned by I770-Cα of UH and Q1010-Cα of CH), and a vector representing the RBD (spanned by Y453-Cα and S399-Cα). The opening of RBD is then defined as the angle (i.e. dot product) between the reference vector and the RBD vector. The twist of RBD is defined as the dihedral angle that is formed by I770-Cα and Q1010-Cα of the reference vector, and S399-Cα and Y453-Cα of the RBD vector.

**Coordinates for NTD rearrangements:** To describe NTD movement, we measure a tilting and torsion of NTD relative to the central helix (CH) (cf. **Fig. 4a**). For this, we introduce a reference vector that represents CH (spanned by A1016-Cα and Q1005-Cα), and a second vector representing the NTD (spanned by L189-Cα and K195-Cα). The tilt of the NTD is then defined as the angle (i.e. dot product) between the reference CH vector and the NTD vector. The torsion of NTD is defined as the dihedral angle that is formed by A1016-Cα and Q1005-Cα of the reference CH vector, and K195-Cα and L189-Cα of the NTD vector.

**Principal Component Analysis:** Principal component analysis (PCA) was performed on the production ensemble combination of G614 - Delta UP and G614 – Delta DOWN runs, to identify dominant modes of structural variations between the variants during MD simulation. The trajectories were aligned by the S2 domain, and the Cα coordinates of the S2 were superimposed with minimal root mean squared deviation. The following calculation was performed on the Cα coordinates of all residues (S1 and S2). A covariance matrix comprised of the following 3 X 3 blocks was obtained:

$$C_{nn'} = \frac{1}{M}\sum_{m=1}^{M} (\vec{r}_{mn} - \langle \vec{r}_n \rangle) \otimes (\vec{r}_{mn'} - \langle \vec{r}_{n'} \rangle)$$

Diagonalizing this covariance matrix gives the eigenvectors describing the directions of maximal fluctuations, each of which translates to a specific conformational change sampled within the ensemble. The eigenvalue of each mode gives the fractional contribution of that mode to the total structural fluctuations. We analyzed the top two modes with the highest eigenvalues. These calculations were performed using the ‘covar’ and ‘anaeig’ commands of GROMACS 5 [23].

### SUPPORTING FIGURES

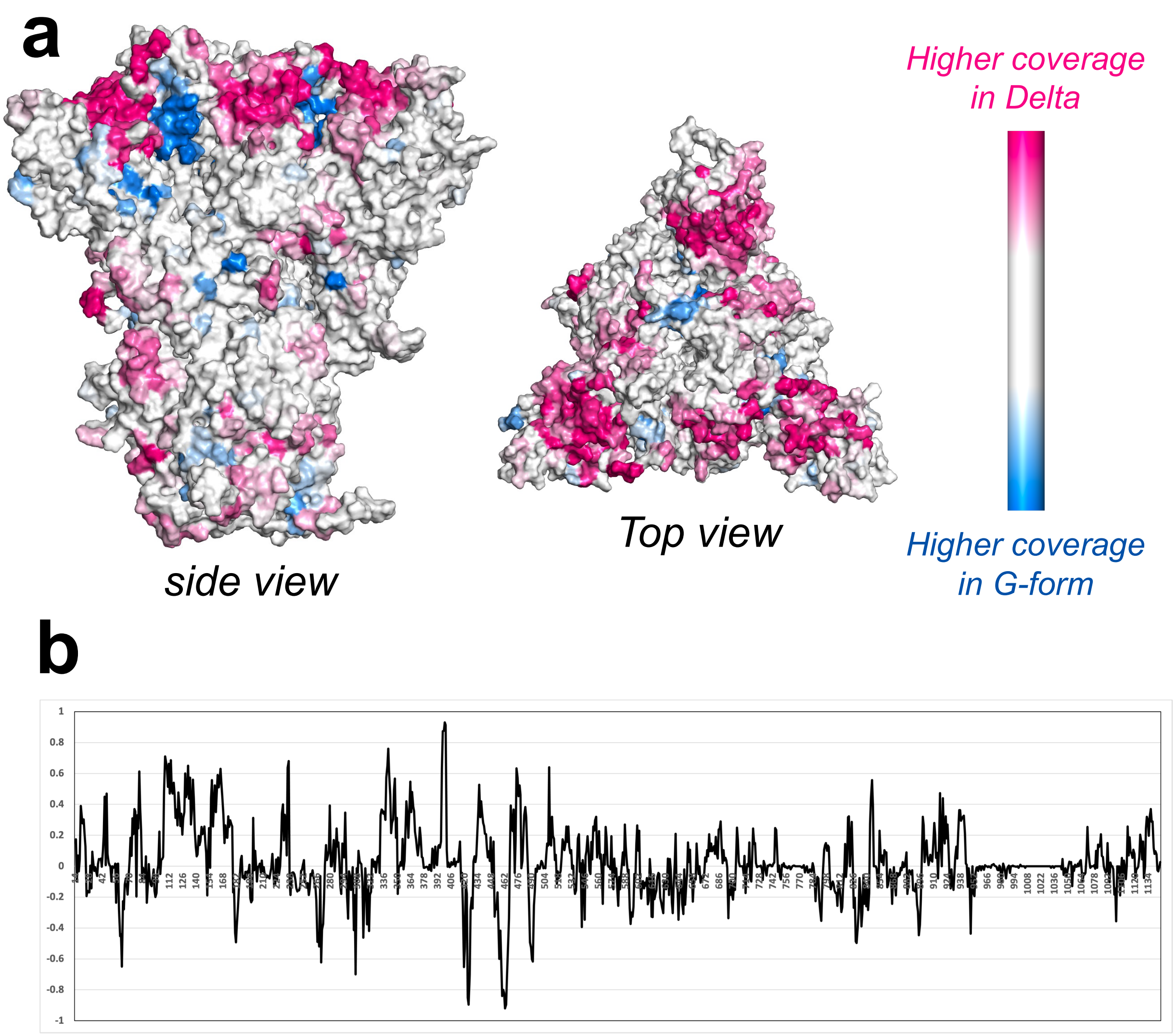


**Figure S1: Differences in glycan coverage between Delta and G-form Spike protein surfaces.** (**a**) Side and top views of the SARS-CoV2 Spike protein. Glycan coverage on the Delta and G-form Spike protein surfaces is measured by the Glycan Encounter Factor (GEF)[19]. GEF difference is represented as a colormap. Dark pink regions represent higher coverage in Delta, and dark blue regions represent higher coverage in G-form. (**b**) Average GEF difference in the RBD-down closed state (Delta minus G-form) plotted as a function of residue number.

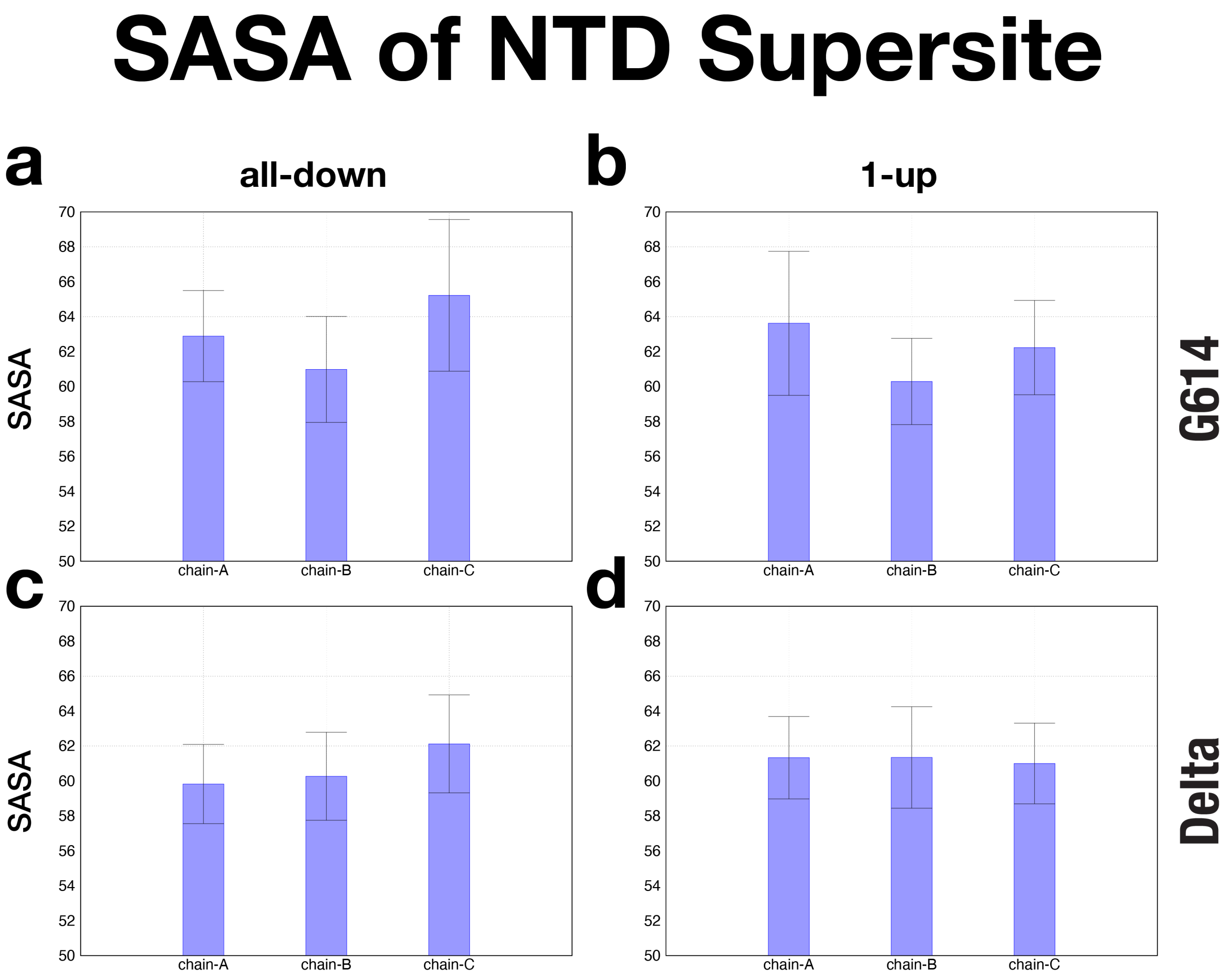


**Figure S2: SASA for the supersite for NTD-binding neutralizing antibodies.** This NTD supersite is defined by loops N1 (14-26), N3 (141-156), and N5 (246-260)[24]. NTD-supersite SASA was calculated for (**a**) G614 "all-down," (**b**) G614 "1-up," (**c**) Delta "all-down," and (**d**) Delta "1-up." In all panels, SASA is shown for chains (or protomers) A, B, and C.

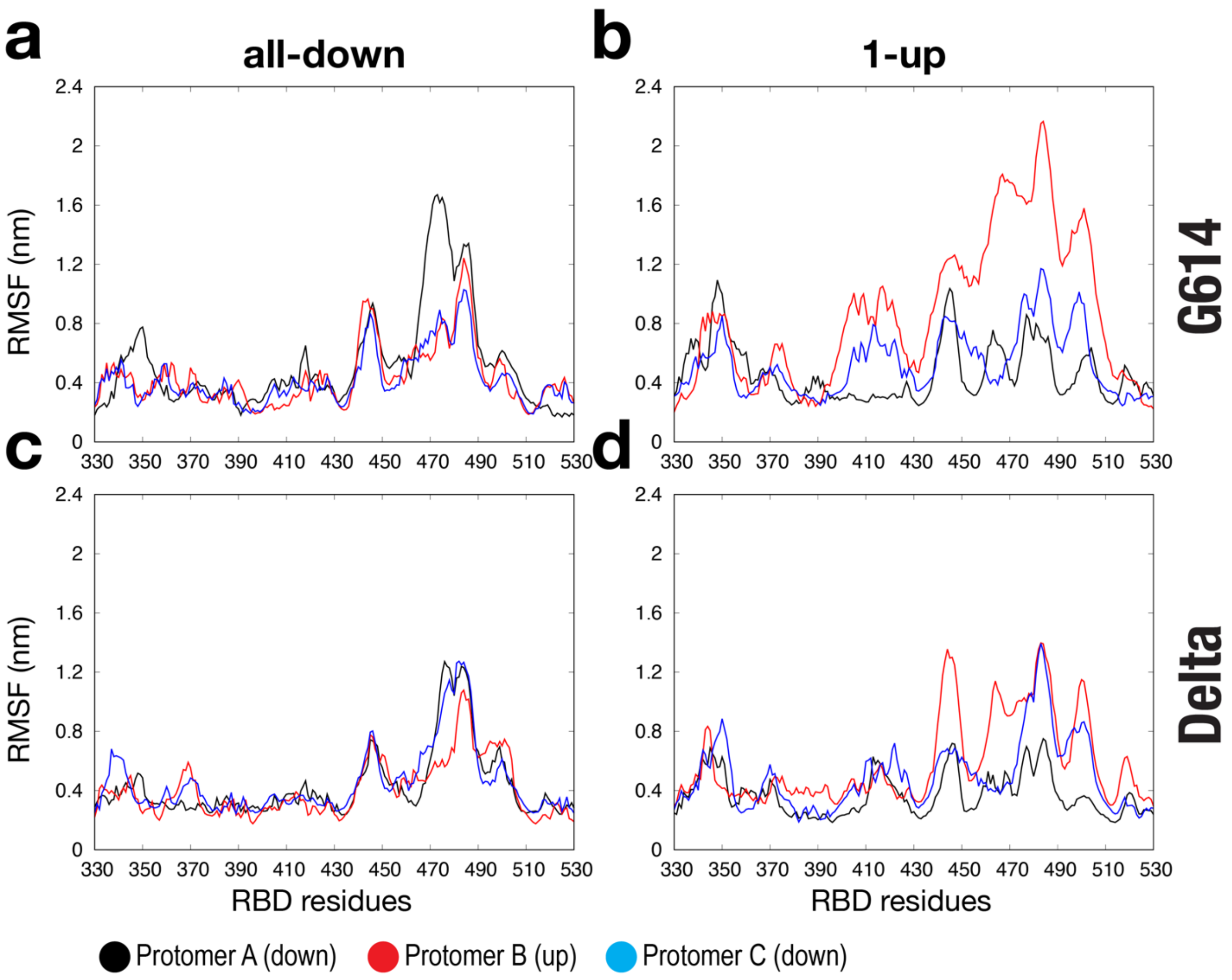


**Figure S3: RMSF as function of RBD residues.** RMSF profiles were calculated for (**a**) G614 “all-down,” (**b**) G614 “1-up,” (**c**) Delta “all-down,” and (**d**) Delta “1-up.” In all panels, RMSF is shown for protomer A (black), B (red), and C (blue). For the 1-up configuration of G614 (cf. panel **b**) or Delta (cf. panel **d**), protomer B has the up-RBD, while protomers A and C have down-RBDs.

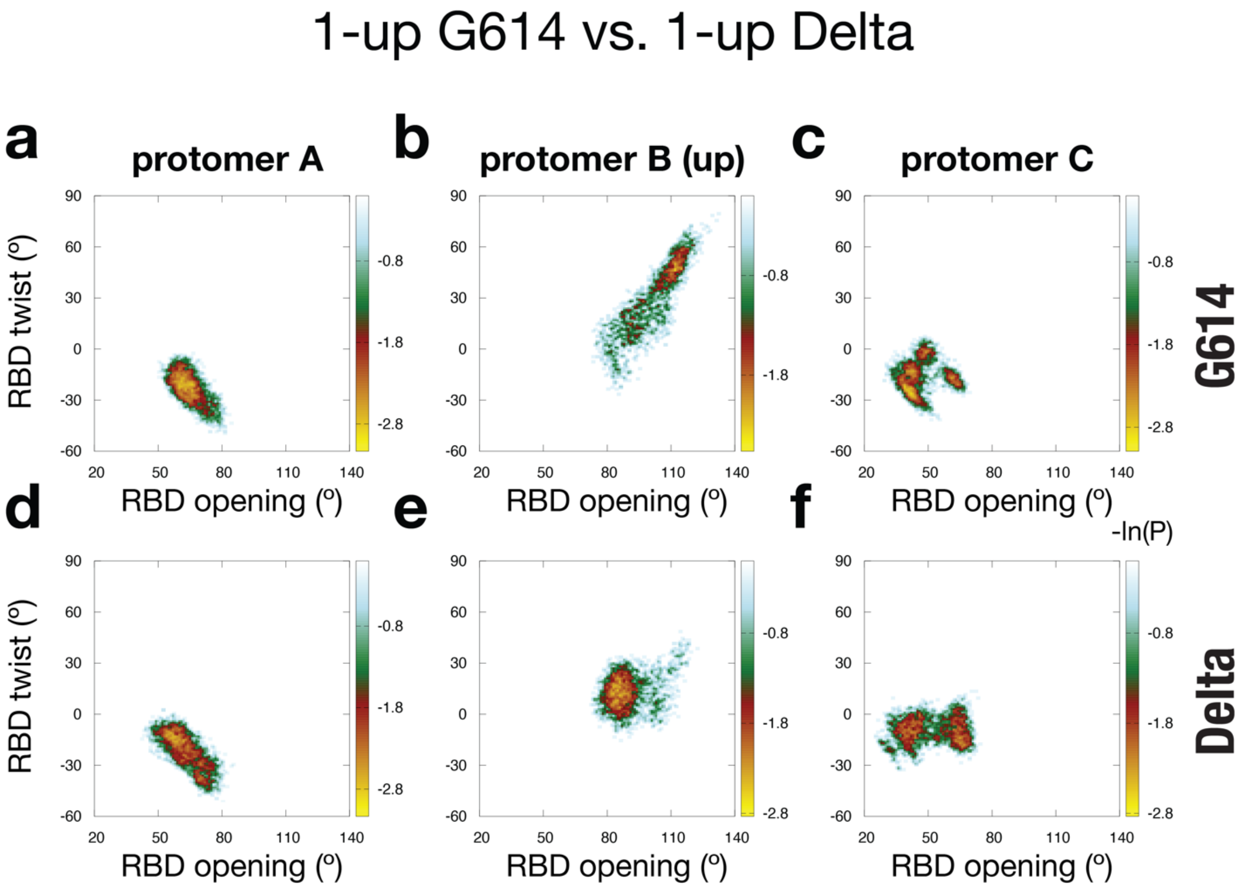


**Figure S4: RBD movement for 1-up G614 versus 1-up Delta.** (**a**/**b**/**c**) 2D distribution along the RBD opening and RBD twist coordinates for each protomer of G614. (**d**/**e**/**f**) 2D distribution along the RBD opening and RBD twist coordinates for each protomer of Delta.

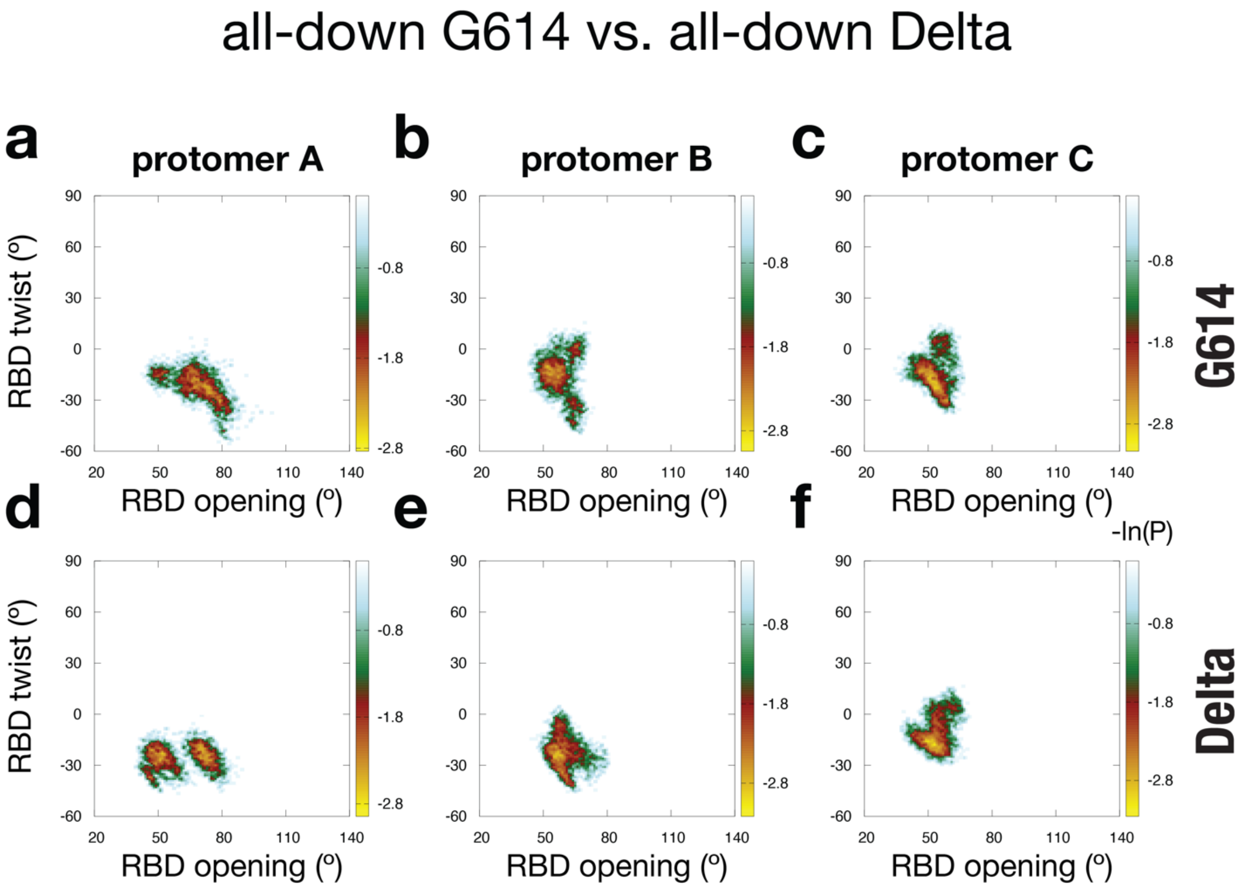


**Figure S5: RBD movement for all-down G614 v. all-down Delta.** (**a**/**b**/**c**) 2D distribution along the RBD opening and RBD twist coordinates for each protomer of G614. (**d**/**e**/**f**) 2D distribution along the RBD opening and RBD twist coordinates for each protomer of Delta.

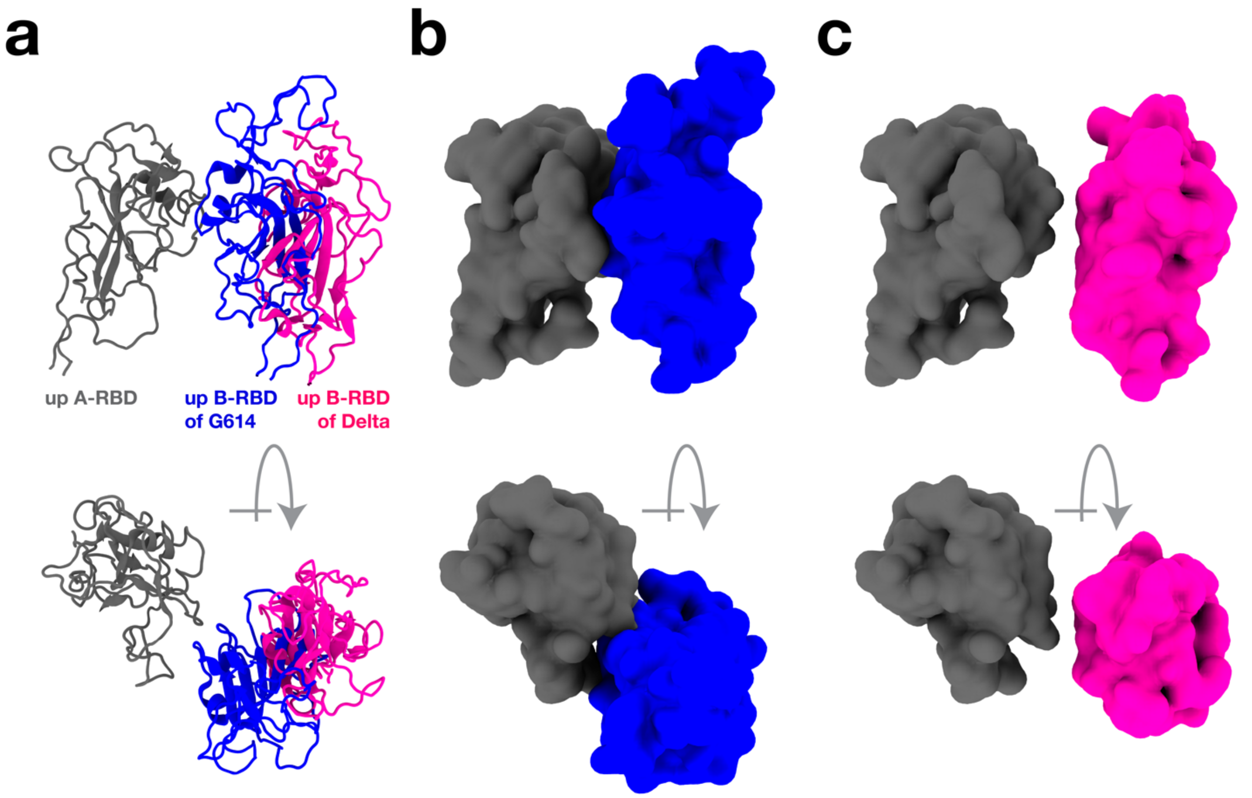


**Figure S6: Distinct up B-RBD conformations in G614 versus Delta.** (**a**) Structural overlay of up B-RBD of G614 (blue) and up B-RBD of Delta (magenta). The G614 RBD (blue) and Delta RBD (magenta) are representative configurations from the "G" and "D" basins as shown in main text **Fig. 3b/c**. To explore how the up B-RBD of each variant may impact the up-transition of the neighboring A-RBD, we modeled an up A-RBD (gray) in protomer A (see main text for details). (**b/c**) To illustrate excluded-volume effects between A- and B-RBDs, panels (**b/c**) show the RBDs (as introduced in panel **a**) in surface representation. (**b**) The orientation of the up B-RBD of G614 (blue) introduces steric hindrance that would impede the up-transition of the neighboring A-RBD (gray). (**c**) In contrast, the distinct orientation of the up B-RBD of Delta (magenta) reduces steric clashes, which would facilitate the up-transition of the A-RBD (gray). All protomer overlays were performed using CA-atoms of the central helix (CH) and HR1.

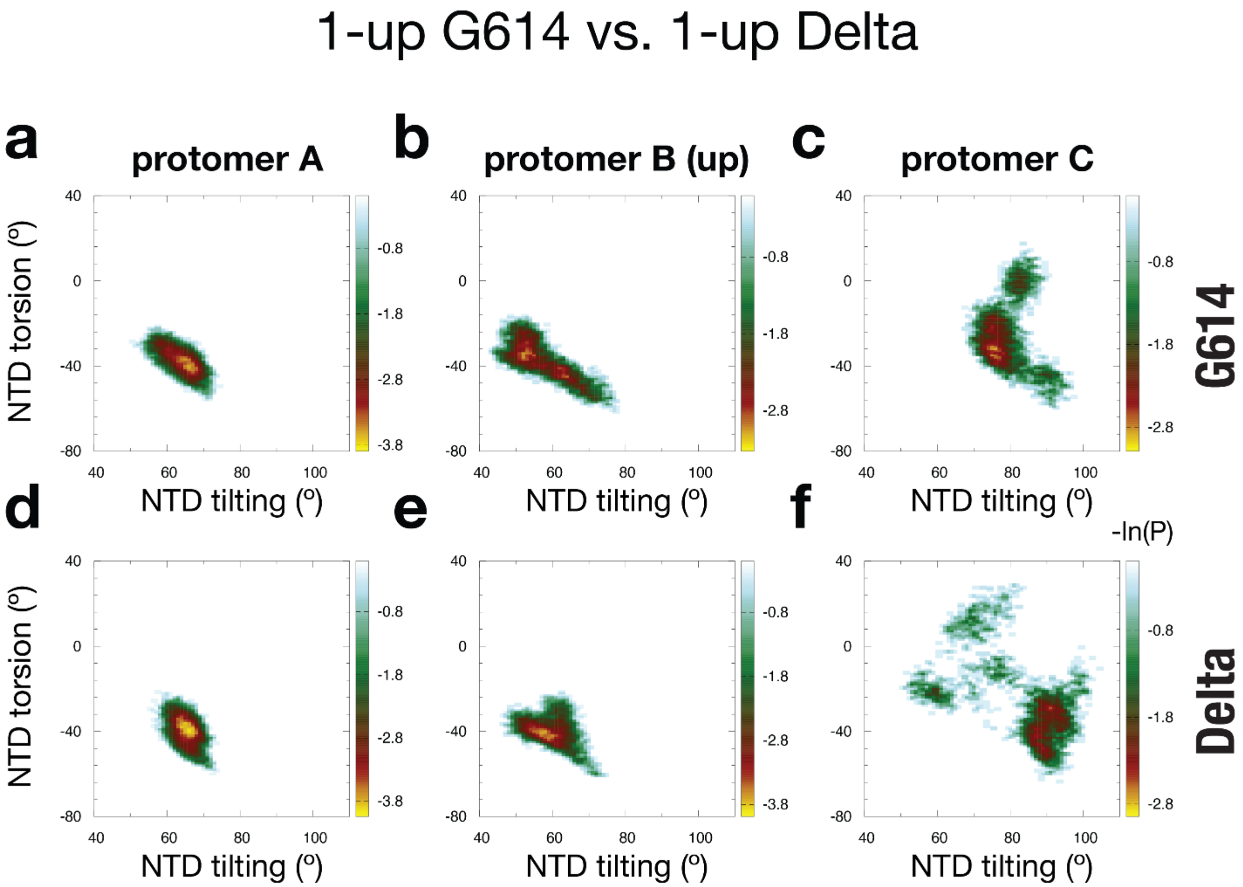


**Figure S7: NTD movement for 1-up G614 versus 1-up Delta.** (**a/b/c**) 2D distribution along the NTD tilting and NTD torsion coordinates for each protomer of G614. (**d/e/f**) 2D distribution along the NTD tilting and NTD torsion coordinates for each protomer of Delta.

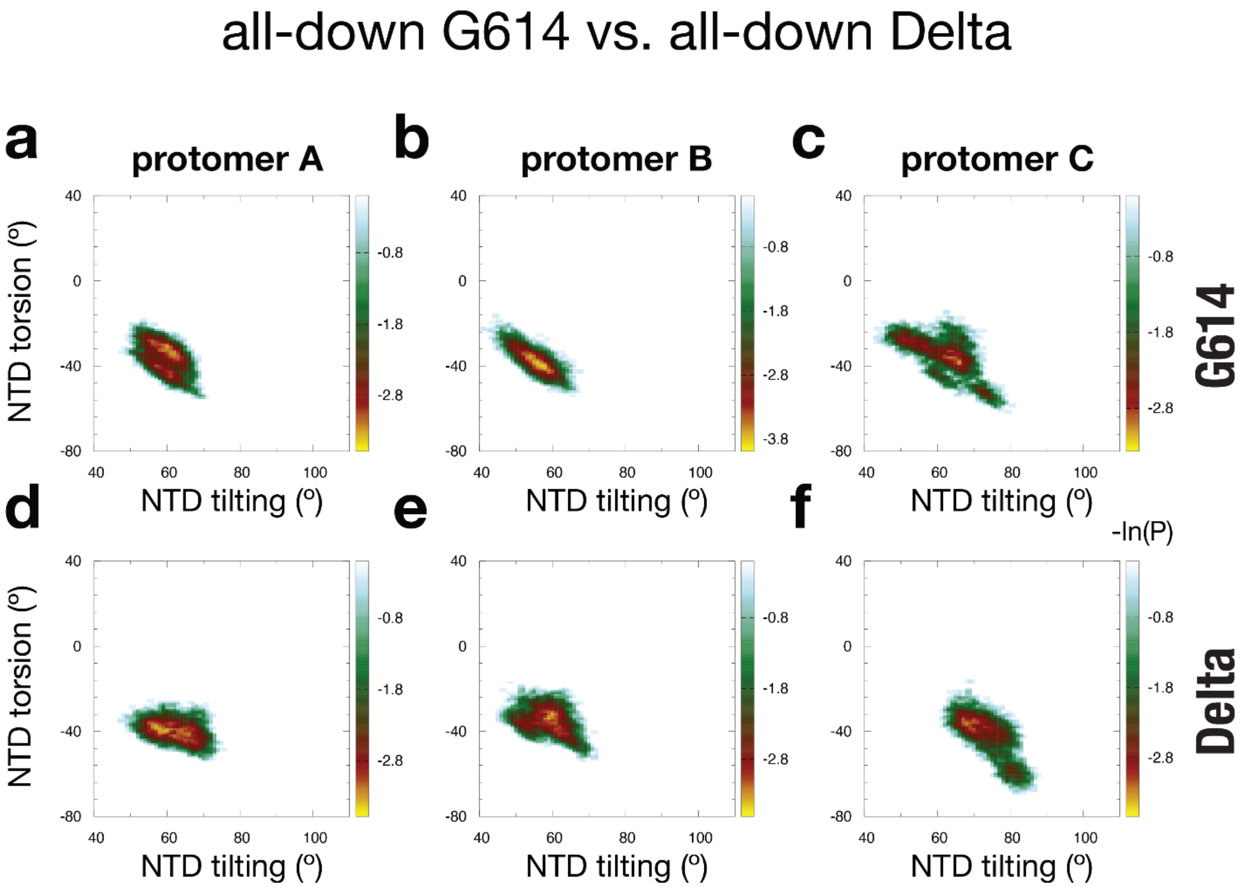


**Figure S8: NTD movement for all-down G614 v. all-down Delta.** (**a**/**b**/**c**) 2D distribution along the NTD tilting and NTD torsion coordinates for each protomer of G614. (**d**/**e**/**f**) 2D distribution along the NTD tilting and NTD torsion coordinates for each protomer of Delta.

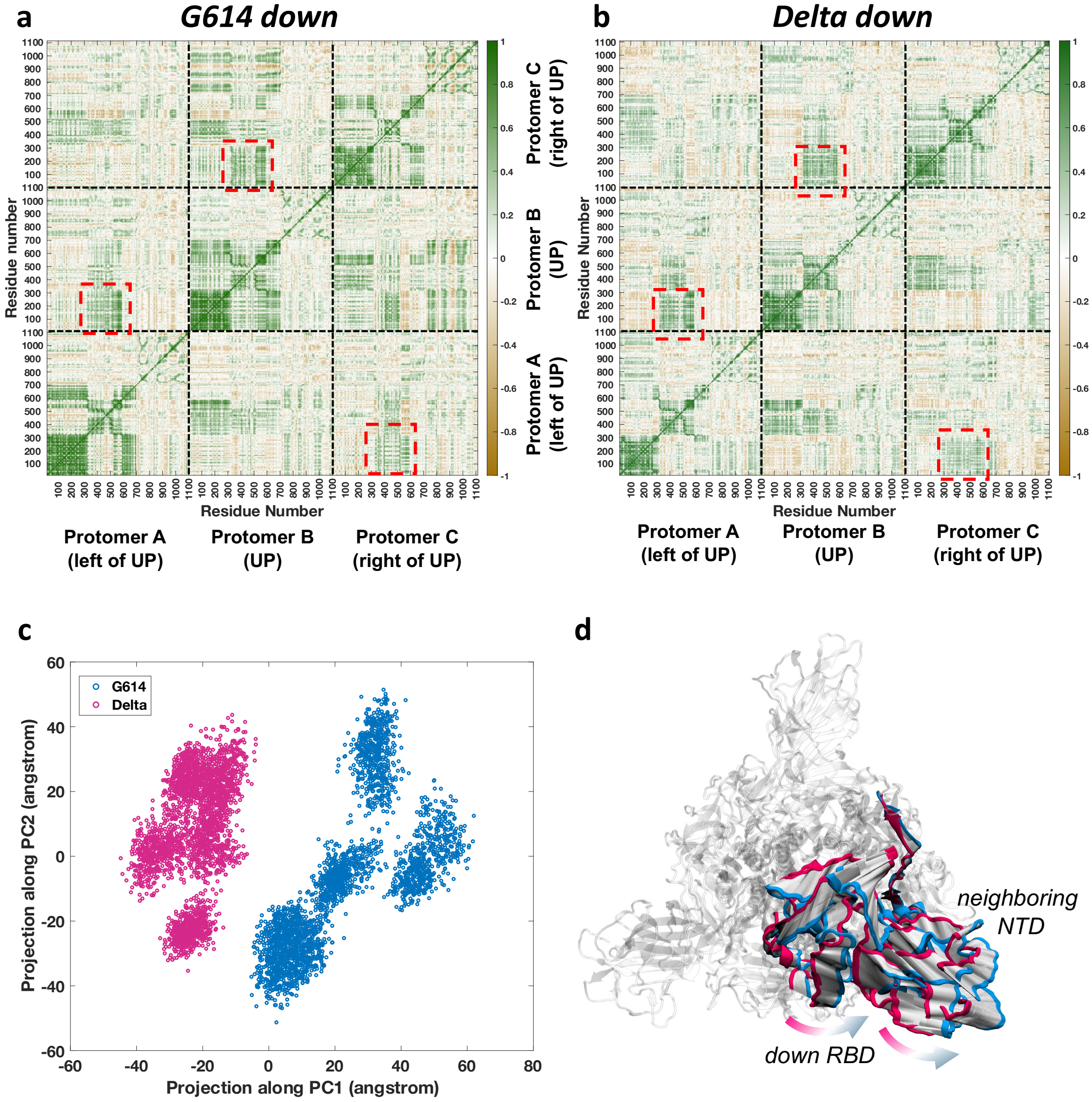


**Figure S9: Differences in correlated motion of RBD between G614 and Delta down conformations.** (**a**) Inter-residue cross-correlation of RMS fluctuations in G614 (all-down), and (**b**) Delta (all-down) conformations. Interprotomer RBD-NTD correlation spaces are indicated by red boxes. (**c**) Projection of the G614 (blue) and Delta (magenta) ensembles along the top two principal component eigenvectors. PC analysis was performed over the two combined ensembles to extract the variances in large-amplitude motions (see methods for details). (**d**) Structural representation of motion along PC1, which describes the dominant difference in motion between G614 and Delta. Magenta structure represents negative PC1 (Delta-like) and blue represents positive PC1 (G614-like), with interpolation given in white. Motion of one RBD (protomer B) and neighboring NTD (protomer C) is shown here.

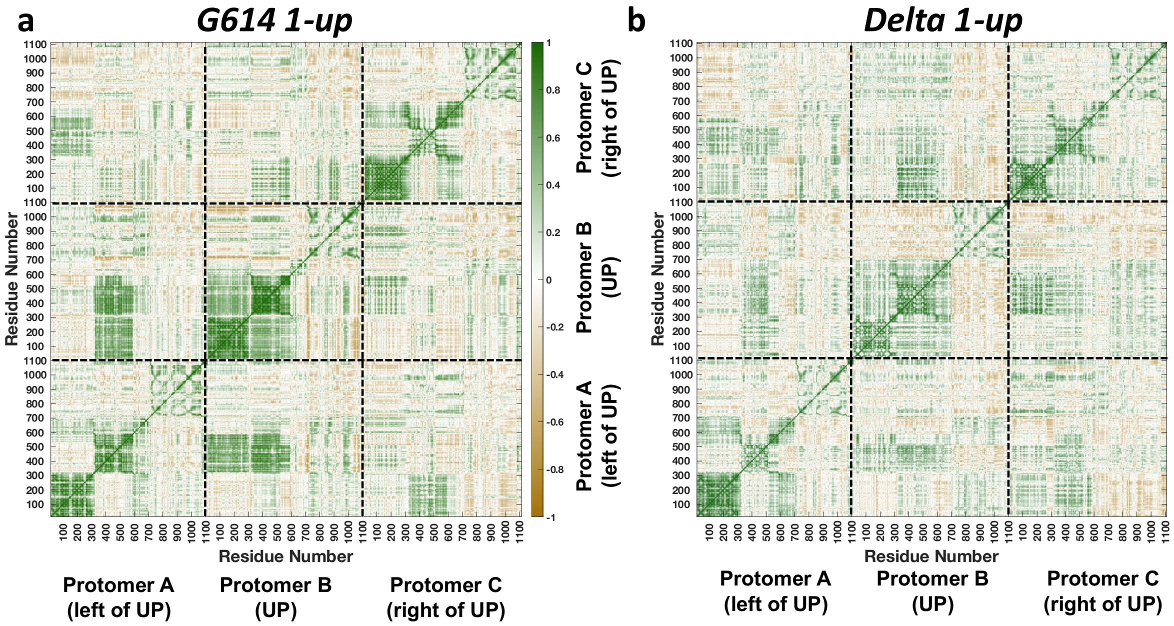


**Figure S10:** Inter-residue cross-correlation of RMS fluctuations in G614 (1-up), and (b) Delta (1-up) Spike protein. In general, the Delta conformation has weaker inter-residue correlations than G614.

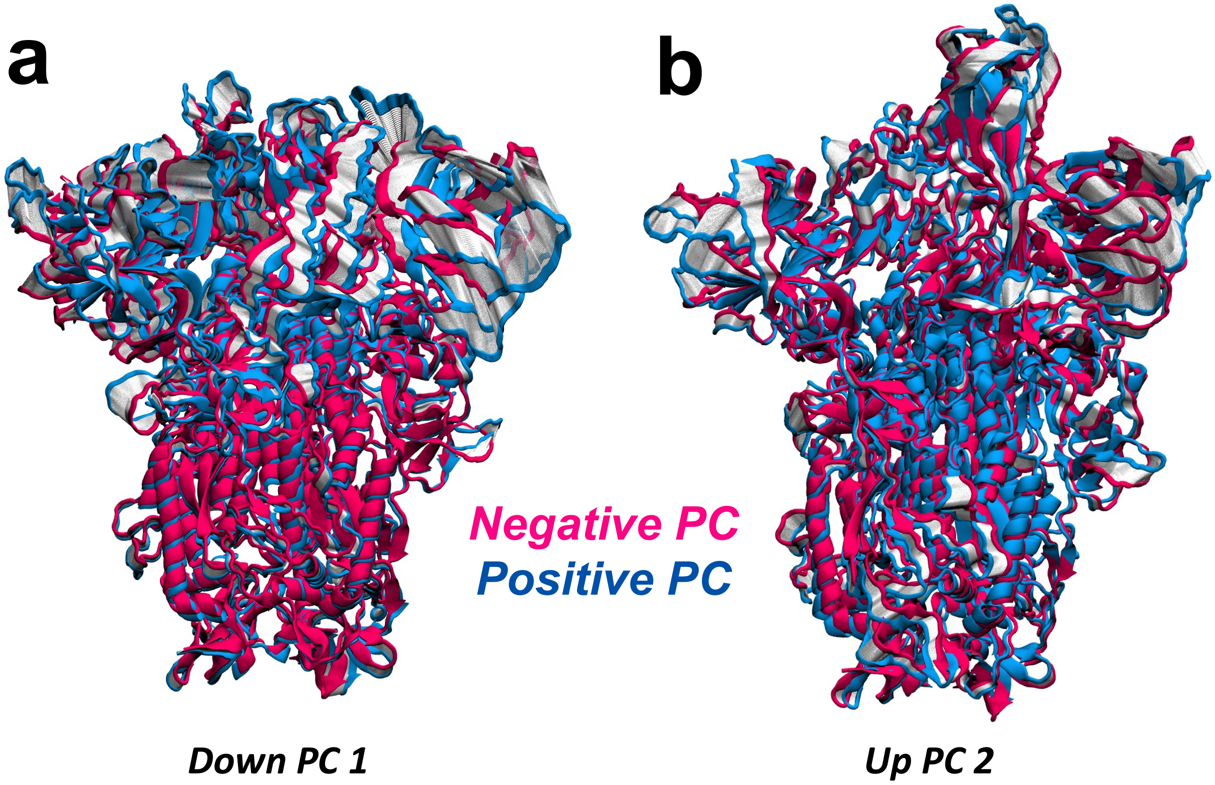


**Figure S11: Principal Component Analysis of cross-correlated fluctuations indicate that dominant motions are mainly confined within the RBD-NTD regions**. (a) PC-1 in RBD-down simulations and (b) PC-2 in 1-RBD up simulations capture the principal differences between G614 and Delta. Magenta structure represents negative PC (Delta-like) and blue represents positive PC (G614-like), with interpolation given in white. Motion along these components are negligible for the central stalk of the Spike.

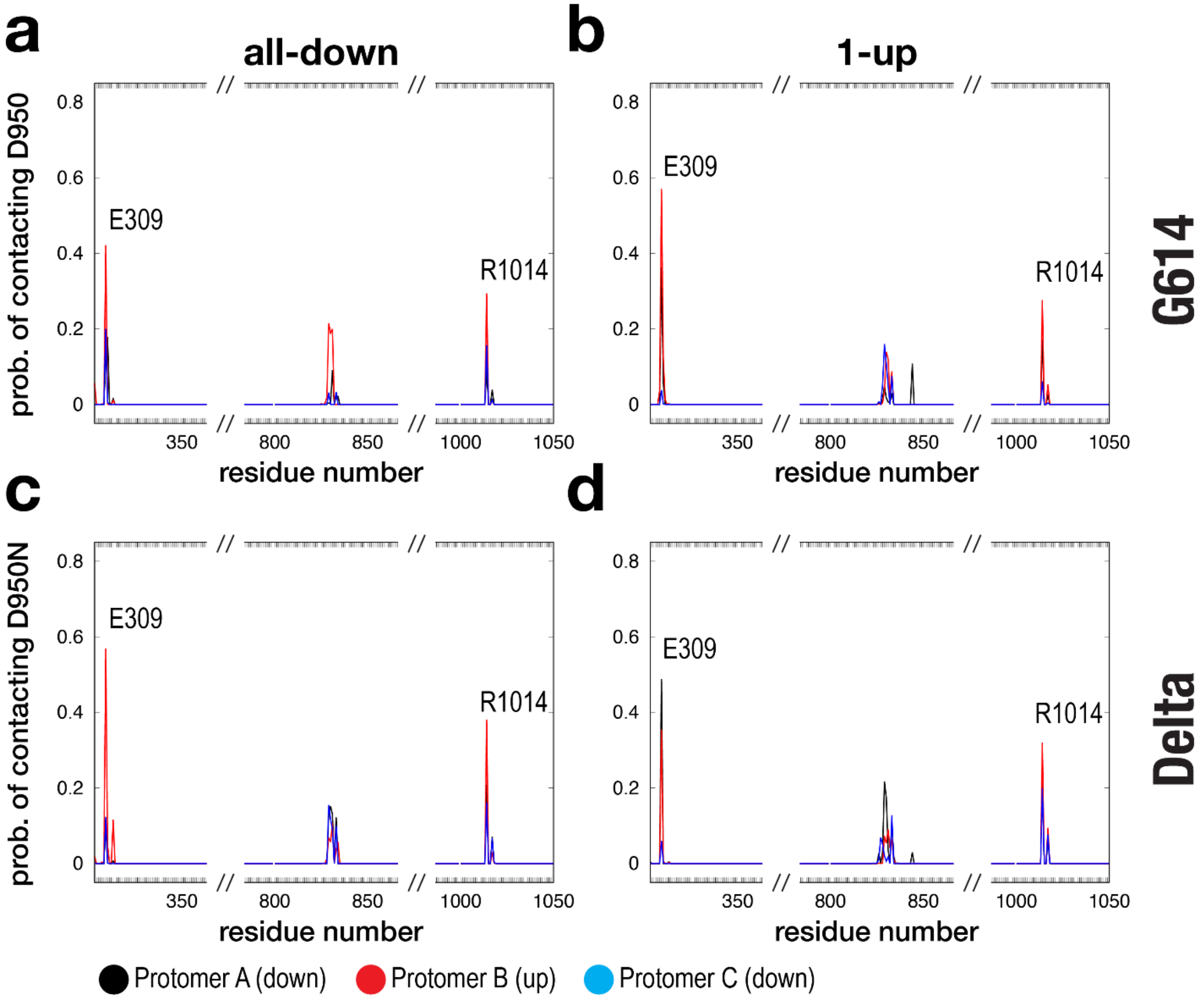


**Figure S12: Interactions with D950N.** To identify residues that form contacts with site 950, we calculated the probability of forming a contact with 950 (y-axis) for any residue (x-axis). Probability profiles are plotted for (**a**) G614 "all-down," (**b**) G614 "1-up," (**c**) Delta "all-down," and (**d**) Delta "1-up." In all panels, profiles are shown for protomer A (black), B (red), and C (blue). For the 1-up configuration of G614 (cf. panel **b**) or Delta (cf. panel **d**), protomer B has the up-RBD, while protomers A and C have down-RBDs.

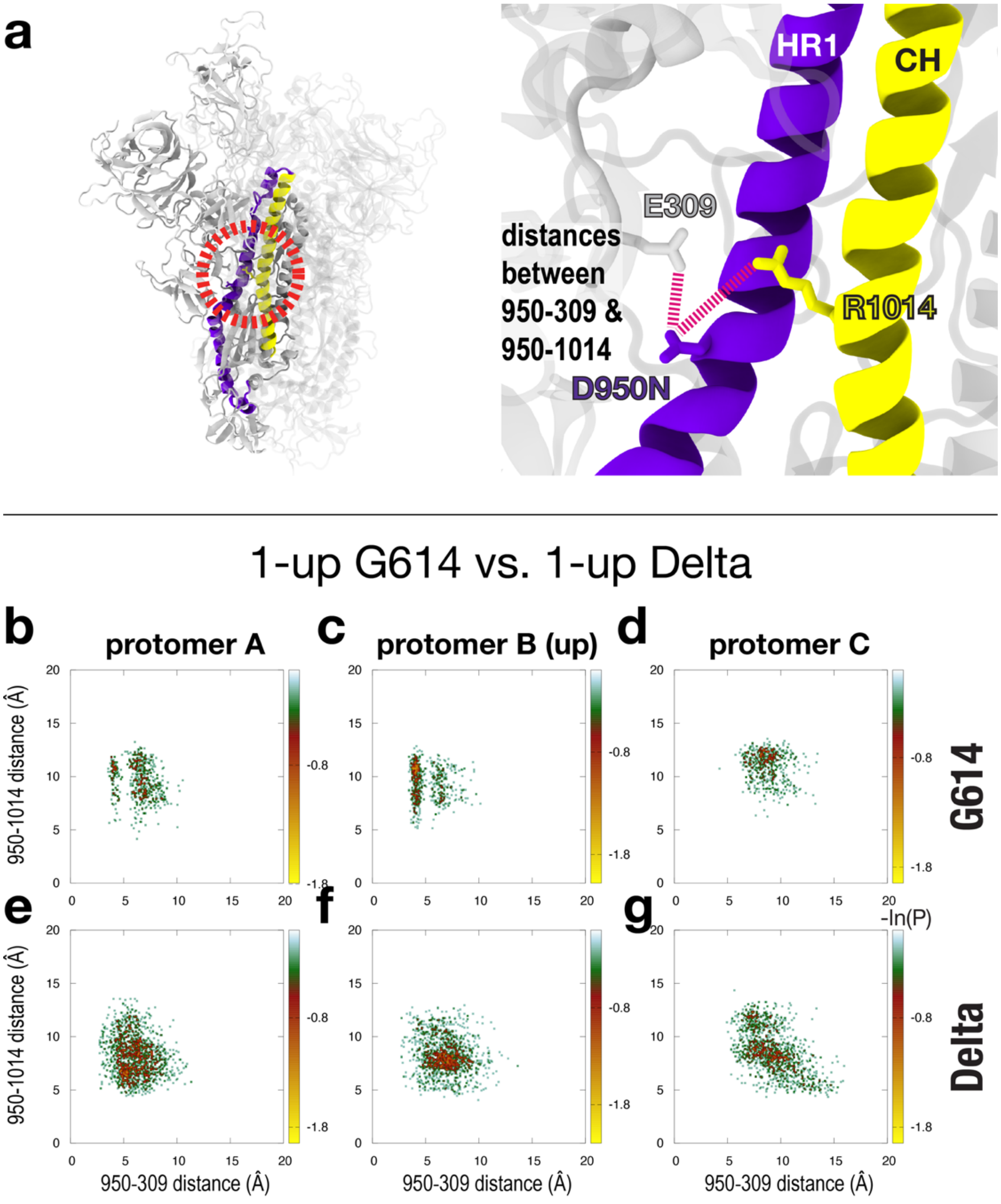


**Caption S13: Sidechain distances to D950N, for 1-up G614 versus 1-up Delta.** (**a**) Structural depiction of distances. The "950-309 distance" is defined as the minimum distance between the E309 O-O group and the D950 O-O group (or N950 N-O) at the end of the sidechains. Similarly, the "950-1014 distance" is defined as the minimum distance between the R1014 N-N group and the D950 O-O group (or N950 N-O) at the end of the sidechains. (**b/c/d**) 2D distribution along the 950-309 and 950-1014 distances for each protomer of G614. (**e/f/g**) 2D distribution along the 950-309 and 950-1014 distances for each protomer of Delta.

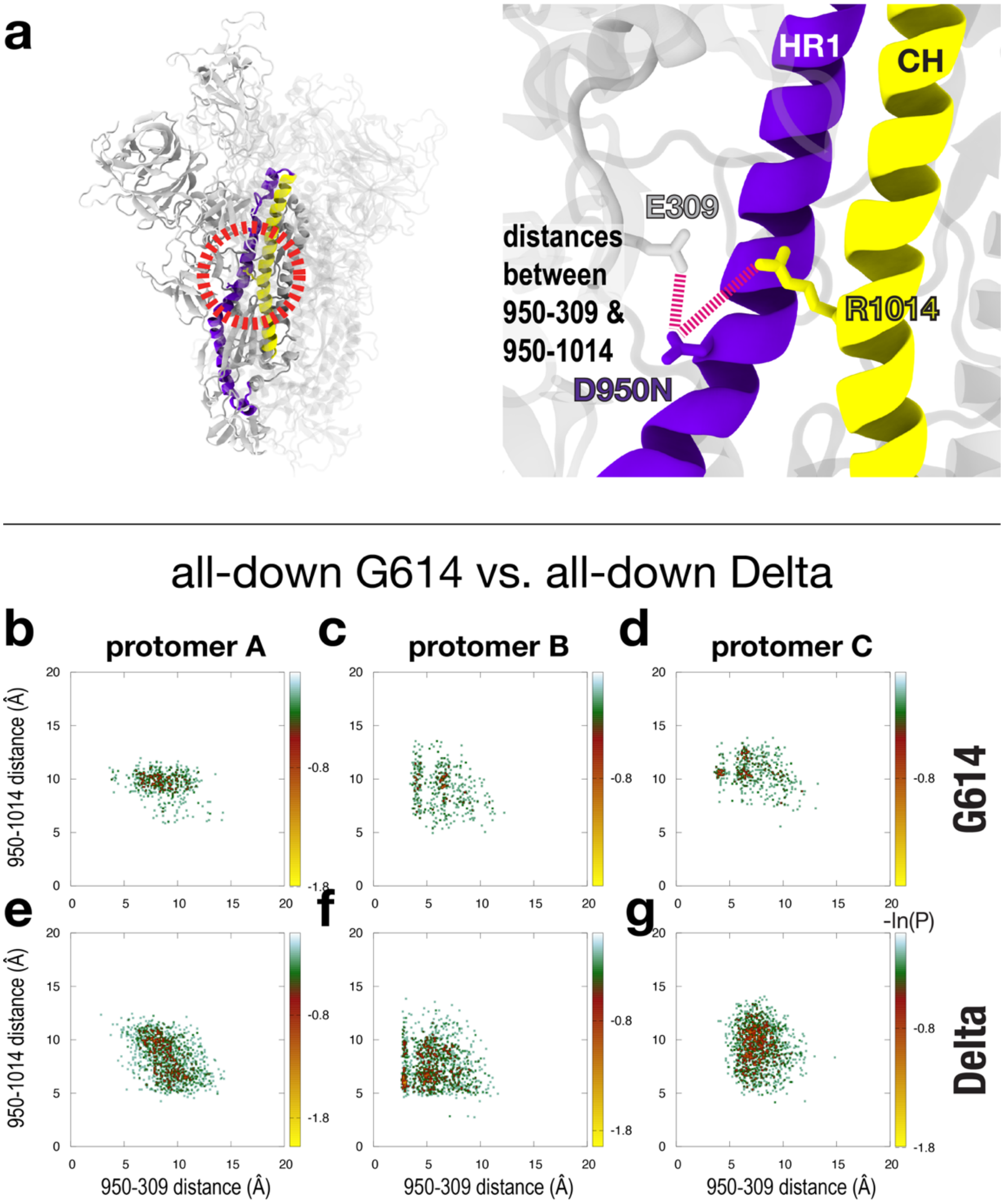


**Figure S14: Sidechain distances to D950N, for all-down G614 v. all-down Delta.** (**a**) Structural depiction of distances. The "950-309 distance" is defined as the minimum distance between the E309 O-O group and the D950 O-O group (or N950 N-O) at the end of the sidechains. Similarly, the "950-1014 distance" is defined as the minimum distance between the R1014 N-N group and the D950 O-O group (or N950 N-O) at the end of the sidechains. (**b/c/d**) 2D distribution along the 950-309 and 950-1014 distances for each protomer of G614. (**e/f/g**) 2D distribution along the 950-309 and 950-1014 distances for each protomer of Delta.

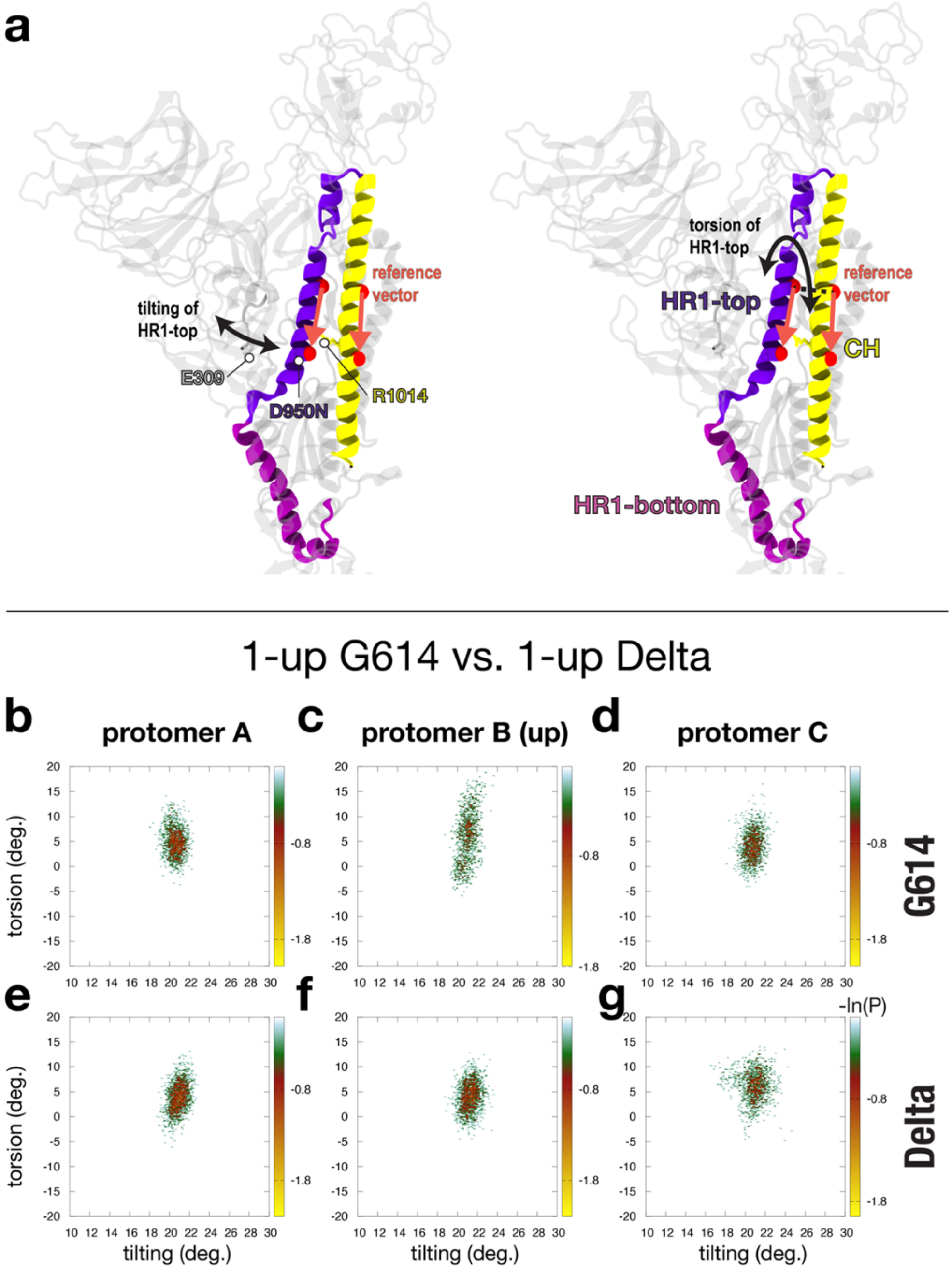


**Figure S15: movement of top region of HR1, for 1-up G614 versus 1-up Delta.** This "HR1-top" region is defined by residues 942 to 986 (violet). (**a**) To describe HR1-top movement, we measure a tilting and torsion of HR1-top relative to the central helix (CH, 987-1035, yellow). For this, we introduce a reference vector that represents CH (spanned by residues A1016 and Q1005), and a second vector representing HR1-top (spanned by residues V951 and L962). The tilt of HR1-top is then defined as the angle (i.e. dot product) between the reference CH vector and the HR1-top vector. The torsion of HR1-top is defined as the dihedral angle that is formed by A1016 and Q1005 of the reference CH vector and L962 and V951 of the HR1-top vector. (**b**/**c**/**d**) 2D distribution along the tilt and torsion angles for each protomer of G614. (**e**/**f**/**g**) 2D distribution along the tilt and torsion angles for each protomer of Delta.

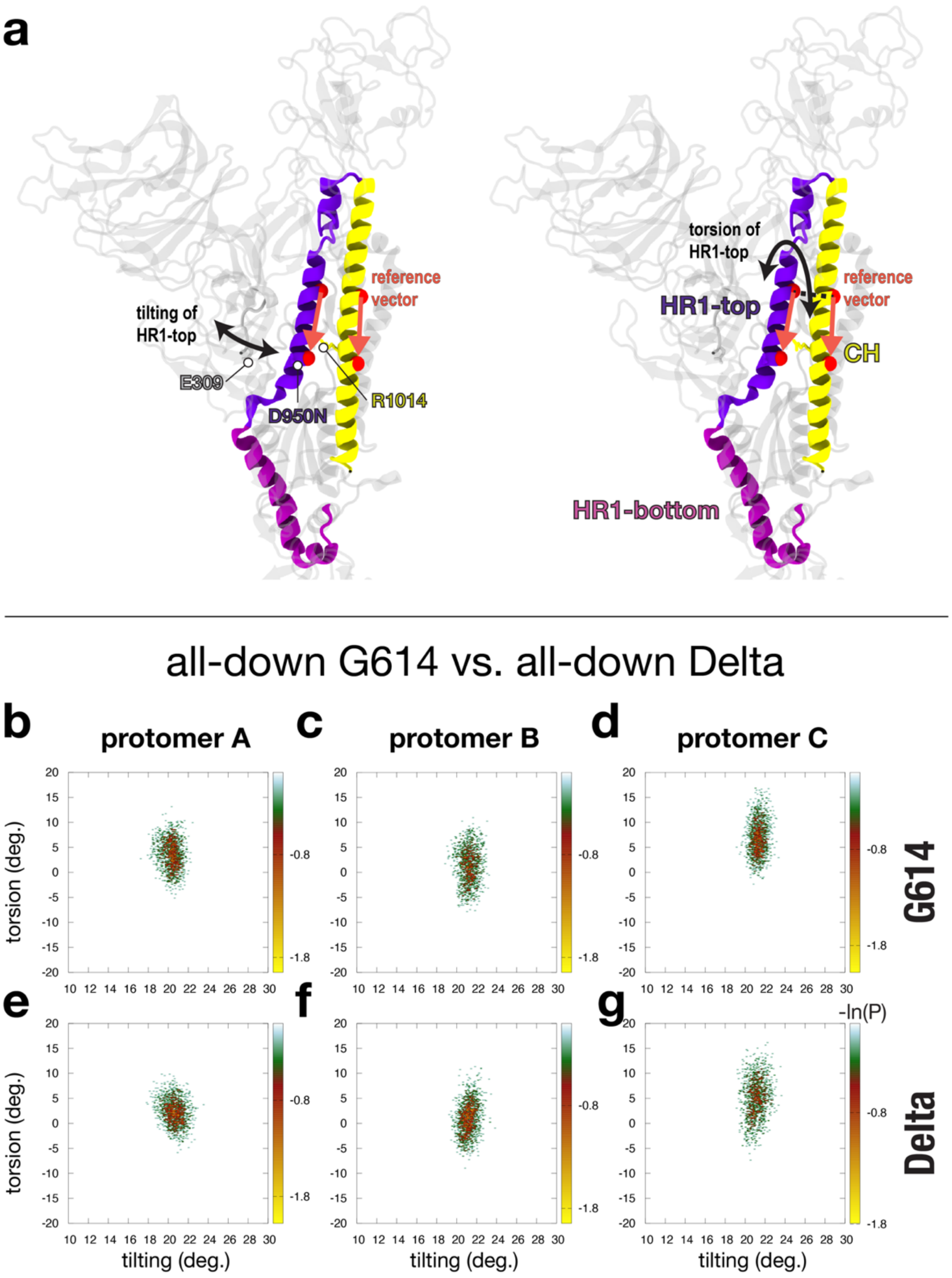


**Figure S16: movement of top region of HR1, for all-down G614 v. all-down Delta.** This "HR1-top" region is defined by residues 942 to 986 (violet). (**a**) To describe HR1-top movement, we measure a tilting and torsion of HR1-top relative to the central helix (CH, 987-1035, yellow). For this, we introduce a reference vector that represents CH (spanned by residues A1016 and Q1005), and a second vector representing HR1-top (spanned by residues V951 and L962). The tilt of HR1-top is then defined as the angle (i.e. dot product) between the reference CH vector and the HR1-top vector. The torsion of HR1-top is defined as the dihedral angle that is formed by A1016 and Q1005 of the reference CH vector and L962 and V951 of the HR1-top vector. (**b/c/d**) 2D distribution along the tilt and torsion angles for each protomer of G614. (**e/f/g**) 2D distribution along the tilt and torsion angles for each protomer of Delta.

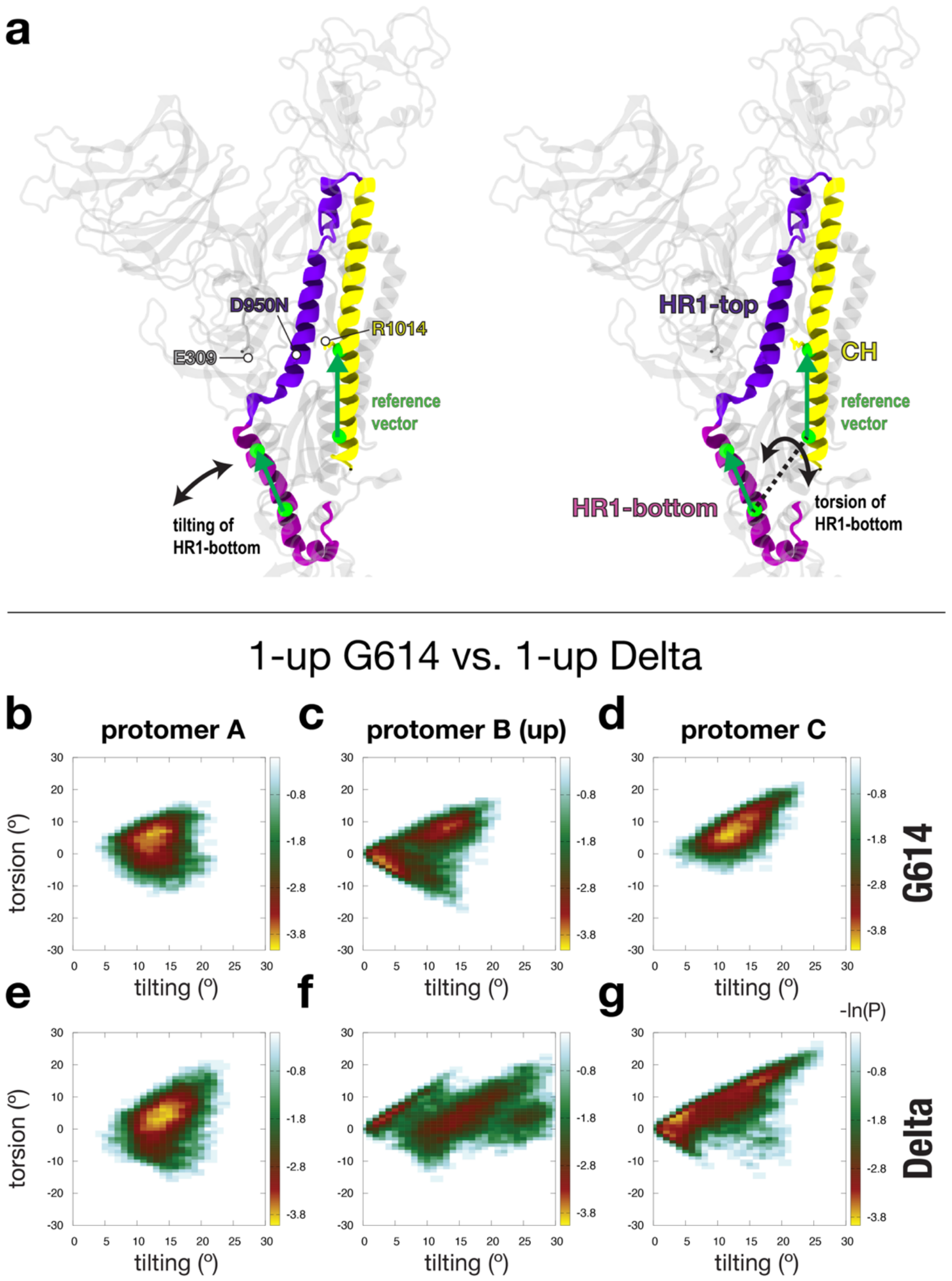


**Figure S17: movement of bottom region of HR1, for 1-up G614 versus 1-up Delta.** This "HR1-bottom" region is defined by residues 910 to 941 (purple). (**a**) To describe HR1-bottom movement, we measure a tilting and torsion of HR1-bottom relative to the central helix (CH, 987-1035, yellow). For this, we introduce a reference vector that represents CH (spanned by residues R1014 and K1028), and a second vector representing HR1-bottom (spanned by residues S937 and Q926). The tilt of HR1-bottom is then defined as the angle (i.e. dot product) between the reference CH vector and the HR1-bottom vector. The torsion of HR1-bottom is defined as the dihedral angle that is formed by R1014 and K1028 of the reference CH vector and Q926 and S937 of the HR1-bottom vector. (**b**/**c**/**d**) 2D distribution along the tilt and torsion angles for each protomer of G614. (**e**/**f**/**g**) 2D distribution along the tilt and torsion angles for each protomer of Delta.

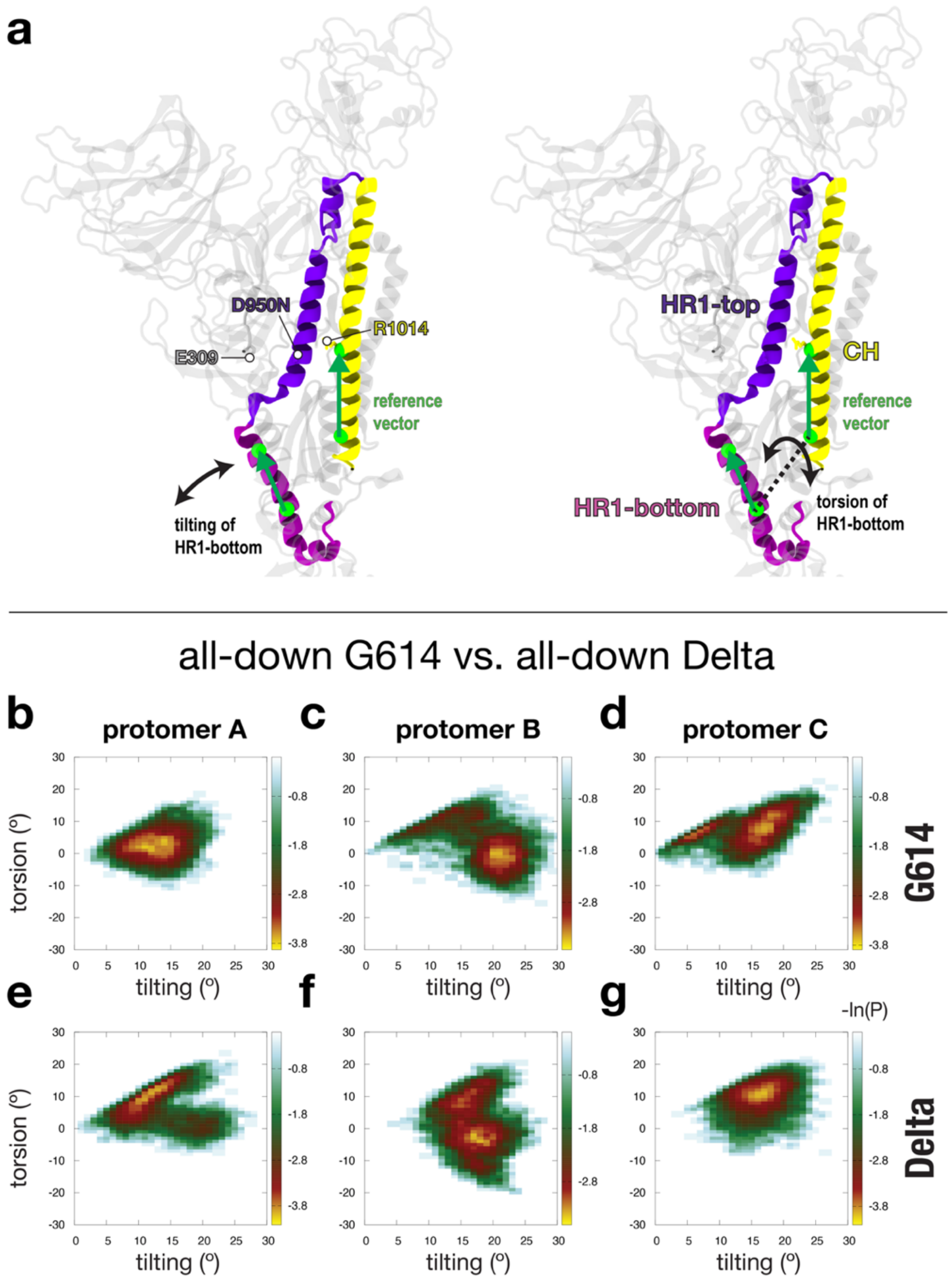


**Figure S18: movement of bottom region of HR1, for all-down G614 v. all-down Delta.** This "HR1-bottom" region is defined by residues 910-941 (purple). (**a**) To describe HR1-bottom movement, we measure a tilting and torsion of HR1-bottom relative to the central helix (CH, 987-1035, yellow). For this, we introduce a reference vector that represents CH (spanned by residues R1014 and K1028), and a second vector representing HR1-bottom (spanned by residues S937 and Q926). The tilt of HR1-bottom is defined as the angle (i.e. dot product) between the reference CH vector and the HR1-bottom vector. The torsion of HR1-bottom is defined as the dihedral angle that is formed by R1014 and K1028 of the reference CH vector and Q926 and S937 of the HR1-bottom vector. (**b**/**c**/**d**) 2D distribution along the tilt and torsion angles for each protomer of G614. (**e**/**f**/**g**) 2D distribution along the tilt and torsion angles for each Delta protomer.

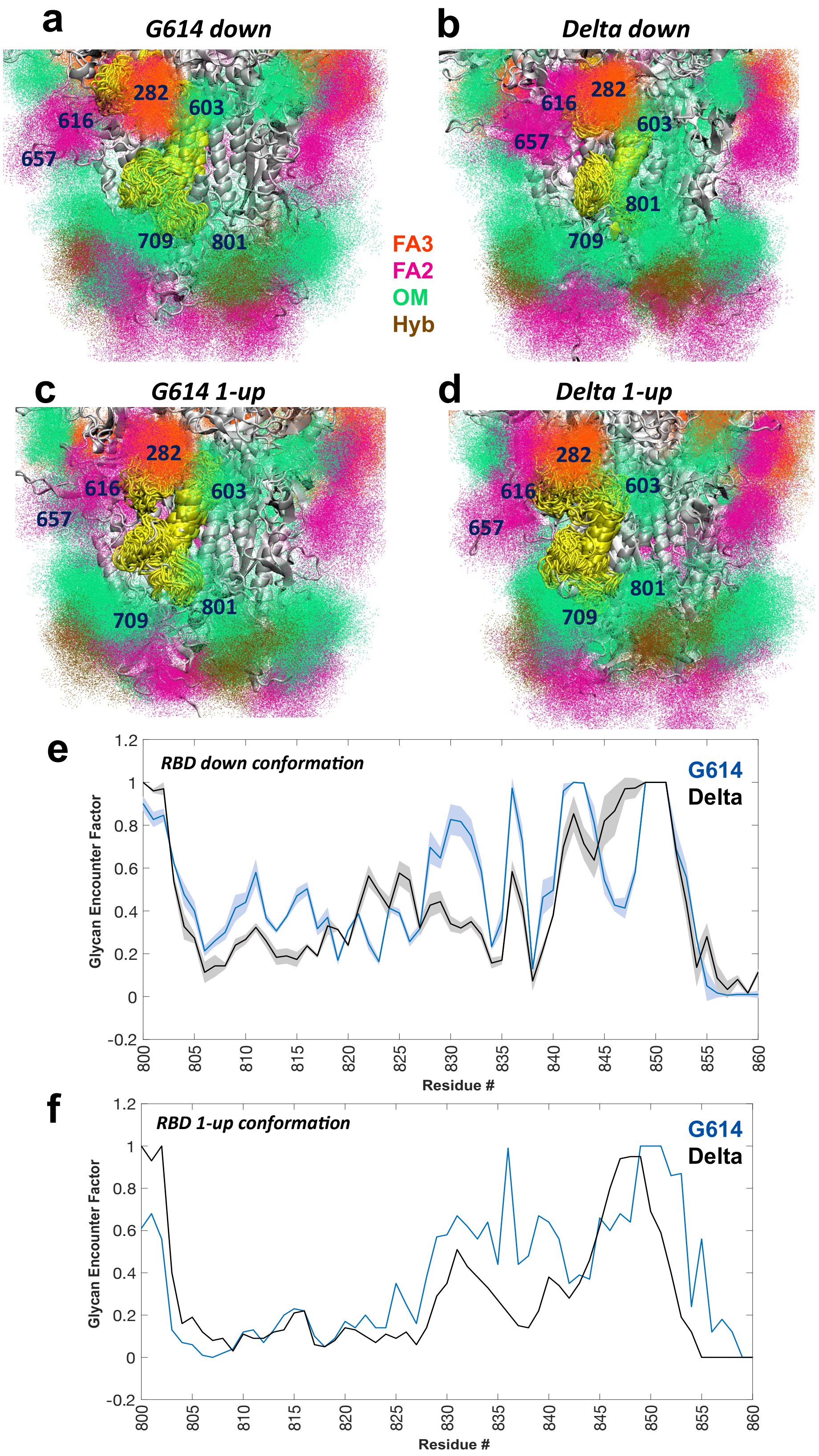


**Figure S19: Glycan shielding of the fusion peptide (residues 800-860) region**. (**a**) G614 down, (**b**) Delta down, (**c**) G614 1-up, and (**d**) Delta 1-up conformations of the fusion peptide region.

Multiple conformations of the fusion peptide residues 800 to 860 are shown in yellow, and the rest of the protein is shown in grey. Glycans proximal to this region are represented as density of points, and colored according to the given key. (**e**) Residue-wise GEF plot for G614 (blue) and Delta (black) Spikes in the RBD down conformation. Mean GEF values from the three protomers are shown, with standard deviation. (**f**) Residue-wise GEF plot for G614 (blue) and Delta (black) Spikes in the RBD 1-up conformation. GEF values only from the up protomer are shown.
